# Mapping interactions within the NMD decapping complex reveals how Upf1 recruits Dcp2 onto mRNA targets

**DOI:** 10.1101/2024.12.18.629111

**Authors:** Nadia Ruiz-Gutierrez, Jeanne Dupas, Elvire Auquier, Irène Barbarin-Bocahu, Claudine Gaudon-Plesse, Cosmin Saveanu, Marc Graille, Hervé Le Hir

## Abstract

Upf1 RNA helicase is a pivotal factor in the conserved nonsense-mediated mRNA decay (NMD) process. Upf1 is responsible for coordinating the recognition of premature termination codons (PTC) in a translation-dependent manner and subsequently triggering mRNA degradation. Multiple factors assist Upf1 during these two consecutive steps. In *Saccharomyces cerevisiae,* Upf2 and Upf3 associated to Upf1 (Upf1-2/3) contribute to PTC recognition but are absent from the Upf1-decapping complex that includes Nmd4, Ebs1, Dcp1, and Dcp2. Despite their importance for NMD, the organization and dynamics of these Upf1-containing complexes remains unclear. Here, we show how distinct domains of Upf1 make direct contacts with Dcp1/Dcp2, Nmd4 and Ebs1 using recombinant proteins. These proteins also bind to each other forming an extended network of interactions within the Upf1-decapping complex. Dcp2 and Upf2 are shown to compete for the same binding site in the Upf1 N-terminal CH domain. This finding accounts for the isolation of two mutually exclusive Upf1-containing complexes from cell lysates. Furthermore, we show how Nmd4-assisted recruitment of Upf1 promotes the anchoring of the decapping protein to the mRNA.

## INTRODUCTION

To ensure correct cellular function, transcriptome homeostasis and attenuate the deleterious effects of gene expression errors, multiple quality control pathways quickly degrade aberrant transcripts. Nonsense-mediated mRNA decay (NMD) is a conserved translation-dependent mRNA degradation pathway that recognizes and degrades mRNAs bearing premature termination codons (PTCs) as well as many physiological transcripts ^1–4^. NMD is implicated in a wide range of cellular processes, including differentiation ^5,6^, stress response ^7^, and immune surveillance ^8,9^. Thus, NMD deregulation is linked to several pathologies, including cancer ^10^, neurodegeneration ^11^, and genetic disorders ^12^.

NMD likely unfolds in two sequential stages: an initial recognition of RNA targets followed by the recruitment of decay factors that irreversibly trigger RNA degradation. These coordinated and successive steps are governed by the evolutionarily conserved RNA helicase Upf1 ^13,14^. Upf1 binds nucleic acids (NA) and is able to translocate directionally along RNA using the energy from ATP hydrolysis ^15–17^. Thus, it can unwind double-stranded NA, translocate along single-stranded NA, and remodel mRNPs *in vitro* ^15,16,18^. Its helicase activity, ATPase activity, and processivity are essential for efficient target recognition and degradation *in vivo* ^17,19^. Together with Upf2 and Upf3, the Upf proteins form a trimeric complex (Upf1-2/3) essential for NMD ^20,21^. However, the precise sequence of events, from Upf1 recruitment to the onset of degradation, is not completely understood.

In the yeast *Saccharomyces cerevisiae* (*S.c.*), NMD is triggered by a PTC distant from the poly(A) tail ^22^. An increased distance between the terminating ribosome and the poly(A) binding proteins decreases the efficiency of translation termination ^23,24^. Consequently, it likely provides time for the Upf1-2/3 trimeric complex to be recruited onto the target mRNA. In mammals and other metazoans, additional layers of regulation contribute to the complexity of NMD ^25^. These include Upf1 phosphorylation ^26,27^ and interactions with specific partners, such as Smg1, Smg6 and Smg5/7 ^28–30^. The presence of a stop codon at least 50-55 nucleotides upstream a spliced junction carrying an exon junction complex elicits NMD ^31–33^.

Degradation of metazoan NMD targets can be achieved by different RNA-decay mechanisms including decapping ^34,35^, endonucleolytic cleavage ^36–38^, and deadenylation ^39,40^. In yeast, the degradation of NMD targets is mostly attributed to deadenylation-independent decapping ^41^. The decapping process relies on the catalytic activity of Dcp2, assisted by its essential co-factor Dcp1 ^42,43^. This heterodimeric complex hydrolyses the 5’ cap structure leading to a 5’ phosphorylated RNA fragment that is tailored for subsequent degradation by the processive exonuclease Xrn1 ^44^. Regardless of the organism, initiation of RNA decay is an irreversible step, committing the RNA to degradation. Thus, triggering of decay has to be precise and stringently regulated to avoid off-target degradation. Decapping activity is regulated by an array of additional co-factors conferring target specificity ^42,43^.

Quantitative mass spectrometry analyses of NMD complexes revealed the existence of two distinct and mutually exclusive Upf1-bound complexes in *S.c.*: the conserved Upf1-2/3 core complex and the Upf1-decapping complex that includes the heterodimer Dcp1/Dcp2 and its co-factor Edc3, as well as the NMD co-factors Nmd4 and Ebs1, and the protein kinase Hrr25^45^. The distinct composition of these two complexes suggests two successive steps linked to potentially distinct conformations and activities of Upf1 during NMD. Similarly, these two distinct Upf1-bound complexes suggest direct regulation of RNA degradation by Upf1-mediated recruitment of decay-inducing partners. Interactions between Upf1 and Dcp2 have been previously established by two-hybrid experiments and co-immunoprecipitation studies in yeast and humans ^40,46,47^. However, understanding Upf1 recruitment mechanisms, as well as the intricate chronology of Upf1-bound complexes, is constrained by the inherent dynamism and transient nature of these complexes. As a result, the mechanisms governing the recruitment and the dynamic interplay of components in NMD complexes remain elusive.

Here, we investigate the molecular bases of the formation and function of the Upf1-2/3 and Upf1-decapping complexes in yeast by dissecting direct protein-protein interactions *in vitro* using recombinant proteins. We demonstrate that Upf1 directly interacts with Ebs1, Nmd4 and the decapping heterodimer Dcp1/Dcp2. The interaction between the two redundant Upf1-binding domains (UBDs) of Dcp2 competes with Upf2 for interaction with Upf1. Furthermore, we show that Upf1 is anchored to the mRNA in the presence of Dcp2 and Nmd4, contrary to the Upf1-Upf2 complex. Altogether, our thorough biochemical characterization provides new insights into the successive events leading to mRNA degradation during NMD.

## RESULTS

### Dcp2 directly interacts with Upf1 N-terminal CH domain

*S.c.* Upf1 is a 971 amino acid (aa) protein composed of three main domains (Figure 1A): an N-terminal domain rich in cysteine and histidine (CH, aa 54-220), followed by the SF1 helicase domain (HD, aa 221-851), and a less conserved and poorly structured C-terminal region (Ct, aa 852-971). To study the interaction between Upf1 and the components of the decapping complex, we purified several versions of Upf1 recombinant proteins from *E. coli*, including the full-length (FL) protein Upf1 (Upf1-FL) and truncated versions (Figure 1A). Each version of Upf1 was fused to an N-terminal calmodulin-binding peptide (CBP) used for pull-down assays. The N-terminal domain of *S.c.* Dcp2 (970 aa) bears the decapping Nudix domain flanked by an N-terminal regulatory domain (NRD) required for Dcp1 recruitment ^48^ and a short domain that mediates the interaction with its activator Edc3 ^49,50^ (Figure 1B). Within the long unstructured C-terminal domain reside several direct or indirect binding sites for decapping regulatory factors including Edc3, Pat1 and two potential Upf1 binding domains (UBDs) ^47,51,52^. These domains were mapped between aa 434-508 for UBD1 and between aa 687-720 for UBD2 (Figure 1B). We purified a short glutathione-S-transferase (GST)-tagged version of Dcp2 encompassing both UBDs (GST-Dcp2-UBD, aa 434-720) and a longer His-tagged version (Dcp2-L, aa 1-720) that required the co-expression with full-length GST-Dcp1 for efficient purification ^50^ (Figure 1B). For binding assays, we mixed different combinations of recombinant proteins, and interactions were detected by co-purification on calmodulin resin in the presence of Ca^2+^. After extensive washing at high salt concentrations, the eluted proteins were fractionated by SDS-PAGE and directly visualized by Coomassie Blue staining (see Methods).

**Figure 1:**
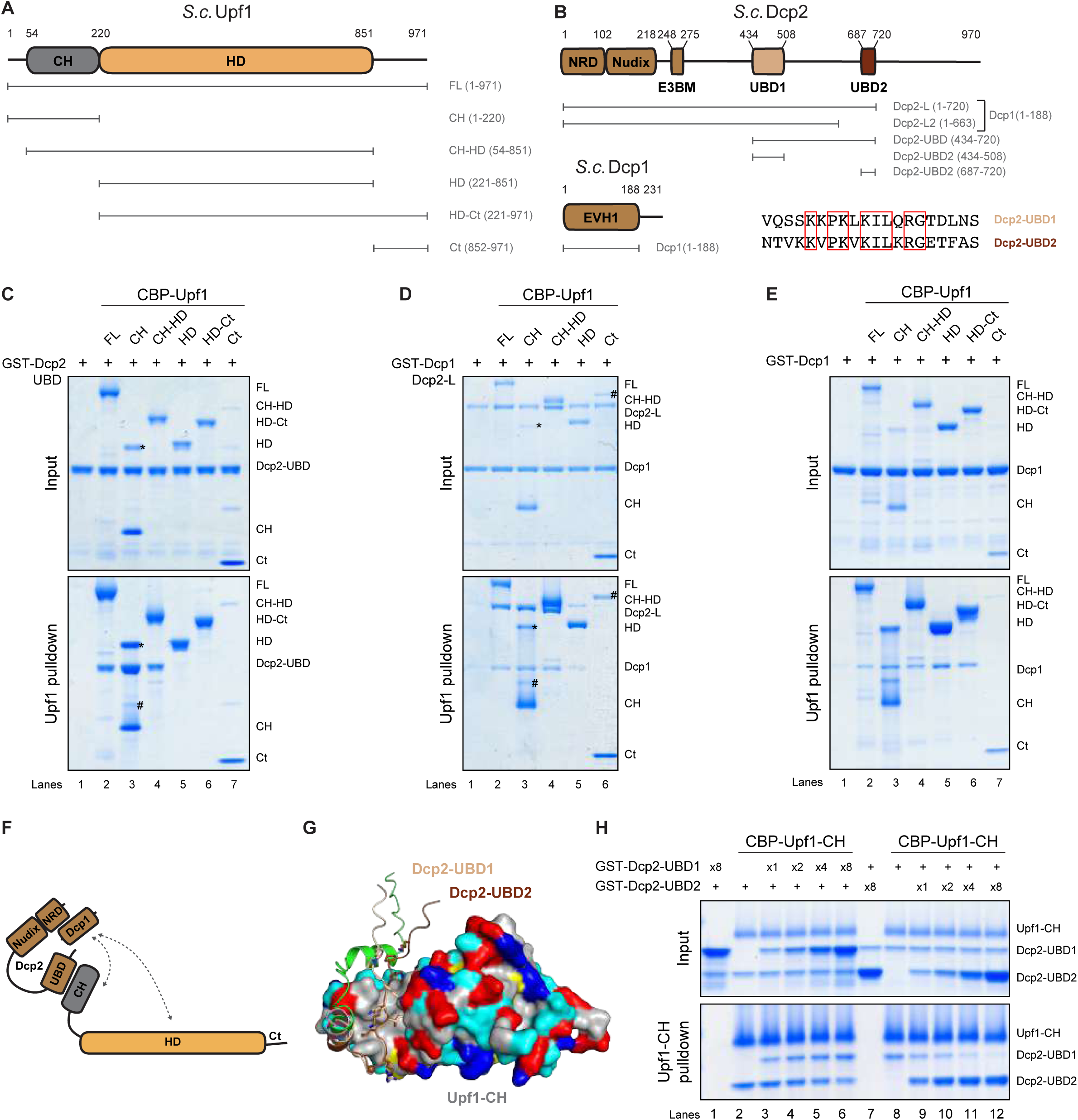
Decapping proteins directly interact with Upf1 CH domain. **(A)** Schematic representation of *S.c.* Upf1 domain arrangement and constructs used for pull-down assays. All constructs feature an N-terminal CBP tag and a C-terminal 6-His tag. Globular domains are depicted as ovals, while lines denote unstructured regions. **(B)** Schematic representation of *S.c.* Dcp2 and Dcp1 constructs used for pulldown assays. Dcp2-UBD (434-720) encompasses both Upf1-binding domains (UBDs) UBD1 and UBD2 whereas isolated Dcp2-UBD1 and UBD2 correspond to truncations 434-508 and 687-720, respectively. All truncations bear a GST tag in N-terminus and 6-His in C-terminus. The heterodimers Dcp1/Dcp2-L and Dcp1/Dcp2-L2 were co-expressed and fused to an N-terminal GST tag on Dcp1 (1-188) and a C-terminal 6-His tag on Dcp2-L (aa 1-720 or aa 1-663). Sequence alignment of UBD1 (wheat) and UBD2 (brown) performed using SEAVIEW ^89^. Conserved residues are indicated by red squares. **(C, D, E)** CBP-pulldown assays using Upf1 truncations against Dcp2-UBD (C), Dcp2-L/Dcp1 (D) and Dcp1 (E). Protein mixtures before (input, 17% of total) or after precipitation (pulldown) were separated on SDS-PAGE and revealed using Coomassie blue. Precipitates were washed with a solution containing 250 mM NaCl, except if noted otherwise. Protein contaminants are indicated with (#). The band indicated with (*) corresponds to a stable dimerization of the isolated CH domain, which has been identified by mass spectrometry and by western blotting. **(F)** Schematic representation of protein-protein interactions between Upf1, Dcp2 and Dcp1. **(G)** Superposition of AlphaFold3 models of the *S.c.* Upf1-CH domain (colored according to amino acid properties) bound to Dcp2 UBD1 (green) and UBD2 (pink). The Dcp2 UBD1 (aa 455-475) and UBD2 (aa 694-713) motifs are colored in wheat and brown, respectively. The side chain of some conserved residues from the UBD1 (wheat) and UBD2 (brown) are shown as sticks. Amino acid physicochemical properties are color coded as follows: hydrophobic residues in gray, polar residues in cyan, positively charged residues in blue, negatively charged residues in red and Cys in yellow. **(H)** CBP-pulldown competition assays using Upf1-CH as bait against Dcp2-UBD2 in excess of Dcp2-UBD1 (left) and Dcp2-UBD1 in excess of Dcp2-UBD2 (right). The excess in input of Dcp2 is marked in the top panel.

We first verified that none of the different CBP-Upf1 constructs interacted with the isolated GST tag protein (Figure S1A). We observed that the isolated Upf1-CH protein formed an SDS-resistant dimer (Figure S1A, lane 3), which persisted and became enriched during the pulldown assay. When CBP-Upf1 proteins were used as bait, Dcp2-UBD was efficiently coprecipitated with all CH-bearing constructs of Upf1 (Figure 1C, lanes 2-4) but not by the constructs lacking the CH domain (Figure 1C, lanes 5-7). Consistently, the heterodimer Dcp1/Dcp2-L was also pulled down by all the CH-bearing Upf1 constructs (Figure 1D, lanes 2-4). When the stringency of the washing conditions was increased, the interaction was maintained (Figure S1B). Surprisingly, a weak but reproducible and stable interaction was detected between the isolated Upf1-HD domain and the Dcp1/Dcp2-L dimer (Figure 1D, lane 5; Figure S1B, lane 5). Thus, we produced a GST-tagged Dcp1 (aa 1-188) isolated protein (Figure 1B) and we tested the interaction with Upf1 constructs by CBP-pulldown. We observed Dcp1 coprecipitation by all CH-containing Upf1 constructs but also by the isolated HD and HD-Ct domains (Figure 1E, lanes 2-6), indicating that Upf1 may also directly contact Dcp1 through its CH and HD domains. This result was verified using the Dcp1/Dcp2-S heterodimer in which Dcp1-FL was co-expressed with a shorter version of Dcp2 (aa 1-315), which does not include the C-terminal extension bearing the UBD domains (Figure S1C, lanes 2-6). Collectively, these results show that Dcp2 directly and stably interacts with the CH domain of Upf1 *via* its C-terminal UBD-containing domain and that its co-factor Dcp1 binds the CH and HD domains of Upf1 (as summarized in Figure 1F).

Sequence alignments of the UBD domains (UBD1 and UBD2) revealed conserved residues between the two UBD sites (Figures 1B, S2A, S2B). Consistently, AlphaFold3 (AF3) structural prediction of the Upf1-CH domain bound to either Dcp2 UBD1 or UBD2 (Figures 1G, S2C, S2D) showed similar binding mode and both presented high ipTM confidence scores (0.82 and 0.73, respectively) (Figures 1G, S2E, S2F). The conserved hydrophobic residues of both UBDs would similarly interact with the hydrophobic pocket within the CH domain (Figure 1G). The predicted involvement of these hydrophobic residues at the interface is further supported by the resistance of the Upf1-Dcp2 interaction to higher salt concentrations (500mM; Figure S1B). Since the predicted structures indicated a potential competition between both UBD domains for Upf1 binding, we purified GST-tagged isolated Dcp2-UBD1 (aa 435-508) and Dcp2-UBD2 (aa 687-720) (Figure 1B) to test their individual binding to Upf1 and potential competition. Binding assays using the different CBP-Upf1 constructs as bait showed that each Dcp2 UBD domain was efficiently co-precipitated by all CH-bearing constructs: Upf1-FL, Upf1-CH-HD and Upf1-CH (Figure S1D, Figure S1E, lanes 2-4).

Given that both UBD domains were predicted to interact similarly with Upf1-CH (Figure 1G), we next conducted competition assays by pulling down on CBP-Upf1-CH from a protein mixture containing either an excess of Dcp2-UBD1 and stable quantities of Dcp2-UBD2 or *vice versa* (Figure 1H). In the presence of an excess of Dcp2-UBD1, the co-precipitation of Dcp2-UBD2 by CBP-Upf1-CH slightly decreased (Figure 1H, lanes 3-6, Figures S1G, S1H). In the reverse case, Dcp2-UBD1 co-precipitation was highly reduced in excess of Dcp2-UBD2 (Figure 1H, lanes 9-12, Figures S1G, S1H). This suggests that both UBD1 and UBD2 bind the same surface of Upf1-CH and are able to displace each other, with UBD2 having a stronger effect. However, when in excess, both UBD domains were bound by Upf1 similarly (Figure S1H). No effect on either Dcp2-UBD1 or Dcp2-UBD2 binding was detected when incubating with increasing quantities of GST-tag protein (Figure S1F). Next, we quantified the affinity between CBP-Upf1-CH and fluorescently labeled peptides corresponding to either Dcp2 UBD1 (aa 456-479) or Dcp2 UBD2 (aa 694-718) by fluorescence anisotropy. We measured the anisotropy difference between each peptide free in solution or incubated with various concentrations of CBP-Upf1-CH to determine Kd values (Figure S1I). This revealed that CBP-Upf1-CH binds UBD1 and UBD2 with Kd of 60.1 µM and 25.1 µM, respectively, corroborating our observations that UBD2 has a stronger effect than UBD1 in our competition assays (Figures 1H, S1G, S1H).

In agreement with the predicted AF3 structures, this demonstrates that both Dcp2 UBD domains compete for the same binding site on Upf1-CH and that UBD2 has a slightly higher relative affinity for Upf1, suggesting that the relatively higher conservation of UBD2 in fungal proteins (Figures 1B, S2A, S2B) is related with its affinity for Upf1.

### Multiple interactions between Upf1-decapping complex components

In addition to the decapping enzyme, the Upf1-decapping complex also contains the NMD-associated factors Nmd4 and Ebs1 and the decapping activator Edc3. *In vivo* co-precipitation experiments showed the importance of HD and Ct domains of Upf1 for the binding of Nmd4 and Ebs1, respectively ^45^. Nmd4 contains an N-terminal PIN domain (aa 1-168) similar to that of the human endonuclease Smg6 and Smg5, followed by a short disordered C-terminal arm ^45,53^ (Figure 2A). The 3D model of Ebs1 predicted by AF3 (Figure S3A) supports a globular architecture for aa 1-591 flanked by a disordered C-terminal region (aa 592-884) predicted with low pLDDT values ^54–56^. The structured 1-591 domain is composed of an N-terminal 14-3- 3 domain similar to that of the human Smg5/Smg7 proteins, associated with a large helical hairpin subdomain, similar to Est1 ^45^. We purified full-length Nmd4 (aa 1-218) fused to an N-terminal twin-strep (TS) tag, and a soluble version of Ebs1 deleted of the last disordered 293 aa (Ebs1, aa 1-591) fused to an N-terminal GST tag (Figure 2A). We first mixed individually Nmd4 and Ebs1 with the different versions of CBP-Upf1. Nmd4 was co-precipitated by all Upf1 versions except Upf1-Ct (Figure S3B) validating that it directly interacts with the helicase domain of Upf1, as previously suggested ^45^. Additionally, Nmd4 also co-precipitated with the CH domain of Upf1 (Figures S3B, lane 3). To what extent the contacts of Nmd4 with both the CH and HD domains contribute to the interaction of Upf1 with Nmd4 remains unknown. Furthermore, we observed a direct interaction of Ebs1 with Upf1 that is mainly mediated by the C-terminal domain of Upf1, as Ebs1 was efficiently precipitated by Upf1-FL and Upf1-Ct (Figure S3C, lanes 2 and 5, Figure 2B for a schematics of observed interactions). We did not include Edc3, as we could not recapitulate stable interactions with other components of the decapping complex in our experimental conditions.

**Figure 2:**
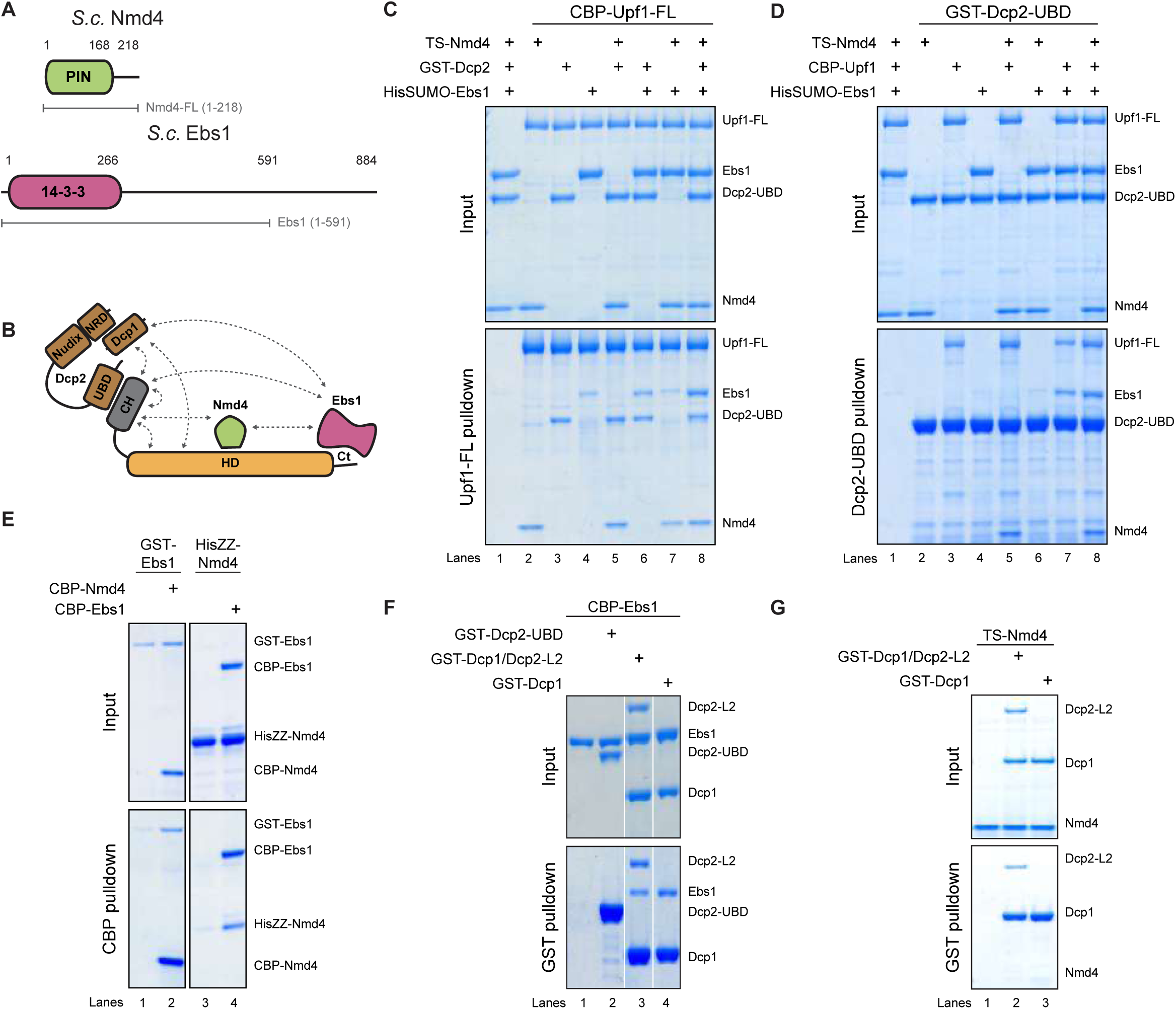
Multiple interactions between Upf1 decapping complex components. **(A)** Schematic representation of *S.c.* Nmd4 and Ebs1 constructs used for pulldown assays. Multiple fusion proteins were produced, with Nmd4-FL (aa 1-218) fused at the N-terminus to either a Twin Strep (TS) tag, CBP, HisZZ, or left untagged. The globular domain of Ebs1 (aa 1-591) was fused at the N-terminus to GST, CBP, or His-SUMO tags. **(B)** Schematic representation of protein-protein interactions between Upf1, Dcp2, Dcp1, Nmd4 and Ebs1. **(C)** CBP-pulldown assays using Upf1-FL against combination of Nmd4-FL, Ebs1(1-591) and Dcp2-UBD as indicated. As previously, protein mixtures before (input, 17% of total) or after precipitation were separated on SDS-PAGE and revealed using Coomassie blue. Protein contaminants are indicated with (#). **(D)** GST-pulldown assays using Dcp2-UBD as bait against combination of Nmd4-FL, Ebs1(1-591) and Upf1-FL as indicated. **(E)** CBP-pulldown assay using CBP-Nmd4 as bait against GST-Ebs1 (left) and CBP-Ebs1 against HisZZ-Nmd4 (right). **(F, G)** GST-pulldown assays using Dcp2-UBD, Dcp1-FL/Dcp2-L2 or isolated Dcp1-FL as bait against CBP-Ebs1 (F) or TS-Nmd4 (G).

We next evaluated whether the individual interactions of Dcp2, Nmd4, and Ebs1 with Upf1-FL interfered with each other. To differentiate each protein and enhance the solubility of Ebs1, we generated HisSUMO-tagged Ebs1 (aa 1-591). CBP-Upf1-FL was mixed with either one partner, a combination of two partners, or all three partners and used as bait (Figure 2C). Upf1-FL precipitated its partners both individually and in combination, indicating no interference among the interactions (Figure 2C, lanes 5-7). To verify our findings, we conducted a similar experiment using GST-Dcp2-UBD as bait (Figure 2D). Dcp2-UBD co-precipitated Upf1-FL but not Nmd4 or Ebs1, whether alone or together (Figure 2D, lanes 2, 4, and 6). In contrast, Dcp2-UBD co-precipitated Nmd4 and/or Ebs1 only in the presence of Upf1-FL (Figure 2D, lanes 5, 7, 8). This demonstrated the successful reconstitution of at least two different trimeric complexes: Upf1-Dcp2-Nmd4 and Upf1-Dcp2-Ebs1 (Figure 2D).

To further investigate the network of interactions among the components of the Upf1-decapping complex we performed GST-pulldowns using the GST-Dcp1/Dcp2-L heterodimer as bait (Figure S3D). Despite the fact that in our experimental conditions, the heterodimer Dcp1/Dcp2 shows a weak stability, we observed that Dcp1/Dcp2 directly interacted with Upf1-FL and Ebs1 individually (Figure S3D, lanes 3 and 4). In agreement with our previous pull-downs, Nmd4 was only co-precipitated in the presence of Upf1 and all three proteins were co-precipitated by GST-Dcp1/Dcp2-L when mixed together (Figure S3D, lane 8). We further explored the interactions between Nmd4, Dcp1/Dcp2 and Ebs1 in the absence of Upf1, the keystone of the complex. We used CBP-Nmd4 or CBP-Ebs1 as bait against GST-Ebs1 and HisZZ-Nmd4 respectively (Figure 2E) and observed a weak but consistent interaction between Nmd4 and Ebs1. Furthermore, we produced a shorter and more stable GST-tagged Dcp2 truncation (aa 1-663) containing UBD1 (Dcp2-L2) (Figure 1B) and used CBP-Ebs1 as bait. We observed a weak but reproducible interaction of Ebs1 with the heterodimer Dcp1/Dcp2-L2 and the isolated Dcp1 protein but not with Dcp2-UBD (Figure 2F) supporting a direct interaction between Ebs1 and Dcp1 and consistent with previous two hybrid results ^57^. This interaction was confirmed by reverse CBP pull-downs with the same proteins (Figure S3E). However, we did not detect direct interactions between Dcp1, Dcp2 and Nmd4 (Figures 2G, S3F). Together, these experiments revealed an intricate network of interactions within the Upf1-decapping complex (summarized in Figure 2B).

### Yeast Upf1-Upf2 interaction is homologous to the human one

The three core NMD factors Upf1, Upf2, and Upf3 form a trimeric complex ^45^ that assembles during, or rapidly after, PTC recognition. The overall organization of the Upf complex appears to be conserved, with Upf2 bridging Upf1 and Upf3 ^20,45,58^. Upf2 is composed of three MIF4G domains, the third of which binds Upf3 ^59–61^, followed by a disordered C-terminal region containing the Upf1 binding domain (aa 933-1089, Figure 3A). In humans, Upf2 has been reported to stimulate the helicase and ATPase activity of Upf1 ^16^. The crystal structure of human Upf1/Upf2 showed that Upf2 binds the CH domain in a bipartite manner, binding one side of the CH domain *via* a β hairpin and the other side *via* an α helix ^62^. The combination of both binding sites renders a highly stable interaction with low dissociation constants ^62^. We modeled the yeast Upf1-CH interaction with Upf2-Ct using AF3 ^63^ (Figure 3B). The model confidently predicted a similar bipartite mode of interaction in which yeast Upf2 would also clamp opposite sides of the CH domain with a β hairpin and an α helix (Figures 3B, S4A). To study the interaction of yeast Upf1 and Upf2 *in vitro*, we purified a truncated version of Upf2 corresponding to the predicted Upf1 binding domain (aa 933-1089, Upf2-Ct) ^16,20^ (Figure 3A). CBP-pulldown assays with the different versions of CBP-Upf1 showed a direct interaction between Upf1-FL, Upf1-CH-HD, Upf1-CH, and Upf2-Ct (Figure 3C), confirming the direct interaction of Upf2 with the Upf1 CH domain. The co-precipitation of Upf2-Ct by CBP-Upf1-FL and CBP-Upf1-CH-HD seemed weaker than the co-precipitation by the isolated CH domain (Figure 3C, compare lanes 2 and 4 to lane 3) as they were slightly above background noise signal. This apparent difference was exacerbated when increasing the stringency of the washes (Figure 3D, lanes 2 and 4), suggesting that the presence of the Upf1 helicase domain interferes with Upf2 binding.

**Figure 3:**
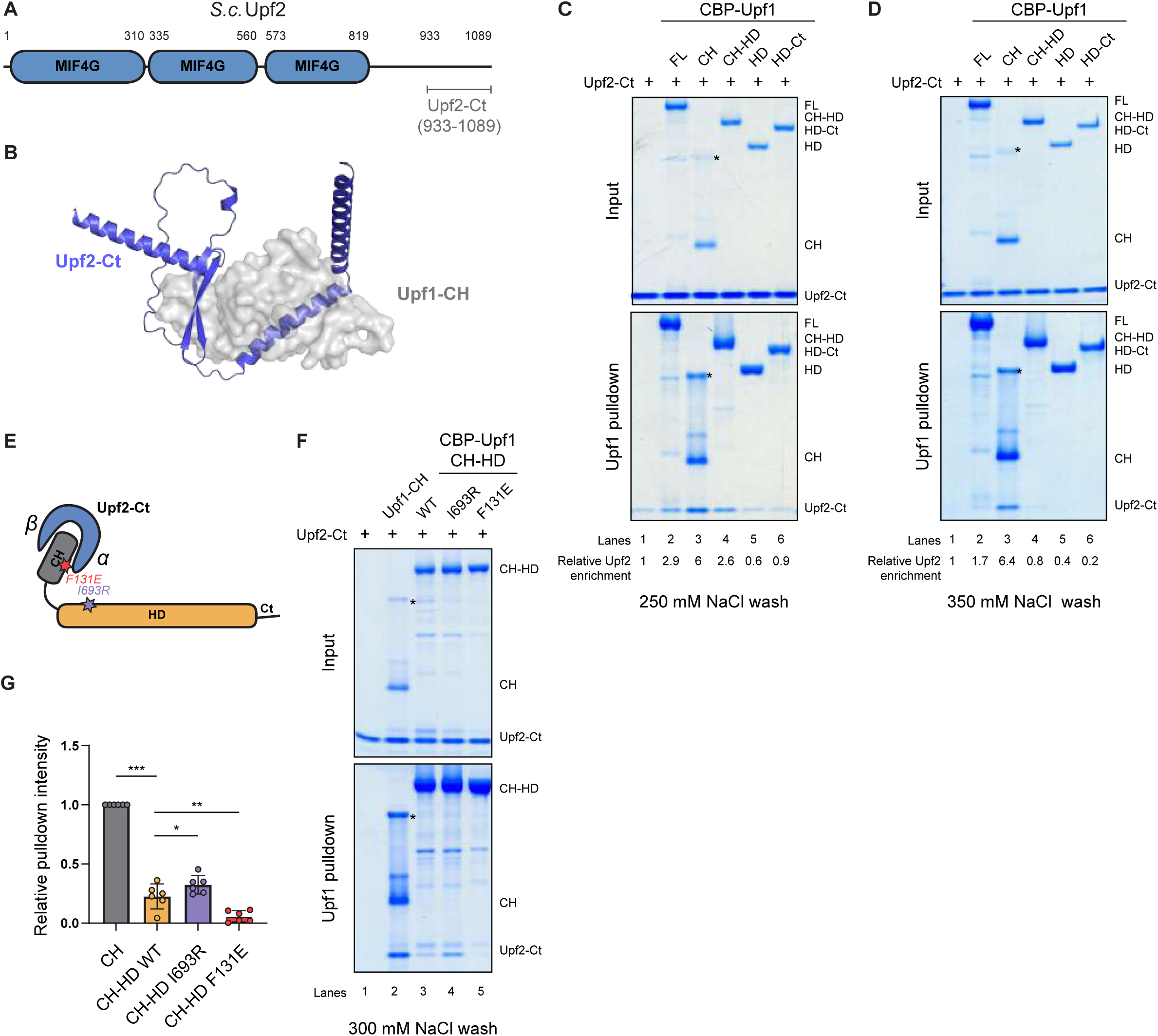
Direct interaction of Upf2 with Upf1 CH domain. **(A)** Schematic representation of *S.c.* Upf2 construct used for pulldown assays. Upf2-Ct (933-1089) is tagged with 6-His in C-terminus. **(B)** Representation of the best AF3 model of the interaction between Upf1-CH domain (grey, surface shown as semi-transparent) and Upf2-Ct domain (blue). **(C, D)** CBP-pulldown assays using Upf1 truncations against Upf2-Ct washed with solution containing either 250 mM NaCl (C) or 350 mM NaCl (D). As previously, protein mixtures before (input, 17% of total) or after precipitation were separated on SDS-PAGE and revealed using Coomassie blue. CH dimer is indicated with (*) and protein contaminants with (#). **(E)** Schematic representation of protein-protein interactions between Upf1 and Upf2. Mutations used in this study of important residues for the intramolecular interaction between CH (red) and HD (purple) of Upf1 are depicted as stars. **(F)** CBP-pulldown assays using Upf1-CH, Upf1-CH-HD WT, Upf1-CH-HD I693R or Upf1-CH-HD F131E against Upf2-Ct. **(G)** Relative band intensity quantification of Upf2 pulldown by Upf1-CH-HD WT, Upf1-CH-HD I693R or Upf1-CH-HD F131E (F), normalized by Upf2 signal intensity pulled down by Upf1-CH (F, lane 2). Error bars denote standard deviations between replicates (n=6). Individual measurements are represented as circles, while the mean of each data series is represented as a column. Unpaired Student t tests were performed to calculate *p* values with a significance threshold set at 0.05 marked with one star. *p* values lower than 0.01 were marked with two stars, and lower than 0.001 were marked with three stars.

To validate the model of Upf1-Upf2 interaction and better understand our results we tested the effect of mutating key amino acid residues. The yeast Upf1-CH-HD protein adopts a closed conformation in which the CH domain is buried within the helicase RecA2 and stalk domains when bound to RNA ^16^. In this structure, the residue I693 in the RecA2 domain interacts with the F131 residue located on the surface of the CH domain (Figure 3E). In humans, the corresponding residue (F192), is involved in the interaction with the α helix of Upf2 ^16^. To investigate whether the yeast interactions followed the same mechanism, we produced CBP-Upf1-CH-HD carrying the I693R or F131E mutations (Figure 3E), which are both expected to prevent the closed conformation of Upf1. The I693R mutation partially restored the co-precipitation of Upf2-Ct by CBP-Upf1-CH-HD under stringent washing conditions (Figure 3F, compare lanes 3 and 4, Figure 3G). As expected, the mutation F131E abolished the interaction with Upf2-Ct (Figure 3F, lane 5, Figure 3G). These results indicate that the same residues are implicated in Upf2 interaction in yeast and humans and confirmed that the intramolecular interaction between the RecA2/Stalk domains of the HD domain and the CH domain of Upf1 hampers binding of Upf2 most likely because only the β hairpin binding site is accessible ^62^.

### Dcp2 and Upf2 compete for interaction with Upf1

Both Upf2 and Dcp2 directly interact with the CH domain of Upf1 (Figures 1 and 3). To explore whether these two Upf1 partners contact the same binding sites, we superposed the AF3 predictions of Upf1-CH/Dcp2-UBD1 and Upf1-CH/Dcp2-UBD2 with the human Upf1-CH/Upf2-Ct crystal structure ^62^ (PDB: 2WJV, Figure 4A). The UBD domains of Dcp2 interact with the outer part of the CH domain in a position similar to that of Upf2’s β hairpin (Figure 4A). This steric clash suggests that Dcp2 and Upf2 cannot simultaneously bind the Upf1 CH domain (Figure 4B). To test this hypothesis biochemically, we conducted competition assays by pulling down CBP-Upf1-CH from a protein mixture containing a fixed amount of Upf2-Ct with gradually increasing quantities of Dcp2-UBD (Figure 4C). The presence of two to eight times more Dcp2-UBD reduced the amount of Upf2 co-precipitated by Upf1-CH (Figure 4C, lanes 2-6, Figure S5A, blue). However, co-precipitation of Dcp2-UBD only marginally increased when used in excess (Figure 4C, lanes 3-6, Figure S5B, brown). The reverse experiment, in which the amount of Dcp2-UBD was constant while the amount of Upf2-UBD increased, led to a comparable reduction of Dcp2-UBD (Figure 4D, lanes 2-6, Figure S5A, brown) but a further increased amount of Upf2-Ct was co-precipitated when in excess (Figure 4D, lanes 3-6, Figure S5B, blue). To confirm the competition between the two partners, we performed sequential binding assays (Figure 4E). We mixed CBP-Upf1-CH with a saturating amount of Dcp2-UBD (Figure 4E, lane 1), washed away the unbound excess of Dcp2-UBD and then added increasing amounts of Upf2-Ct (Figure 4E, lanes 2-5). The latter progressively displaced Dcp2-UBD as previously observed (compare Figure 4D and 4E). In contrast, when Upf1-CH was saturated with Upf2-Ct (Figure 4E, lane 6) before addition of Dcp2-UBD (Figure 4E, lanes 7-10), its displacement was limited (compare Figure 4C and 4E). Our results suggest that Upf2 and Dcp2 compete for the same binding site on the Upf1-CH domain, but that the bipartite interaction of Upf2 with the CH domain offers a more stable grip than that of Dcp2. This was confirmed by anisotropy fluorescence measurements, which allowed to determine a Kd of CBP-Upf1-CH for Upf2-Ct. We first incubated CBP-Upf1-CH with the fluorescently labeled Dcp2 UBD2 (aa 694-718) peptide and then titrated with increasing concentrations of Upf2-Ct. Due to the competition between Upf2-Ct and Dcp2 UBD2 for interaction with CBP-Upf1-CH, the displacement of Dcp2 UBD2 by Upf2-Ct results in a decrease in fluorescence anisotropy, from which we could calculate a Kd value of 0.43 µM for the interaction between CBP-Upf1-CH and Upf2-Ct (Figure 4F). This value is about 140 and 60 times lower than those determined for Dcp2 UBD1 and Dcp2 UBD2 peptides, respectively. This Kd between yeast proteins is very similar to one previously measured between human proteins ^62^.

**Figure 4:**
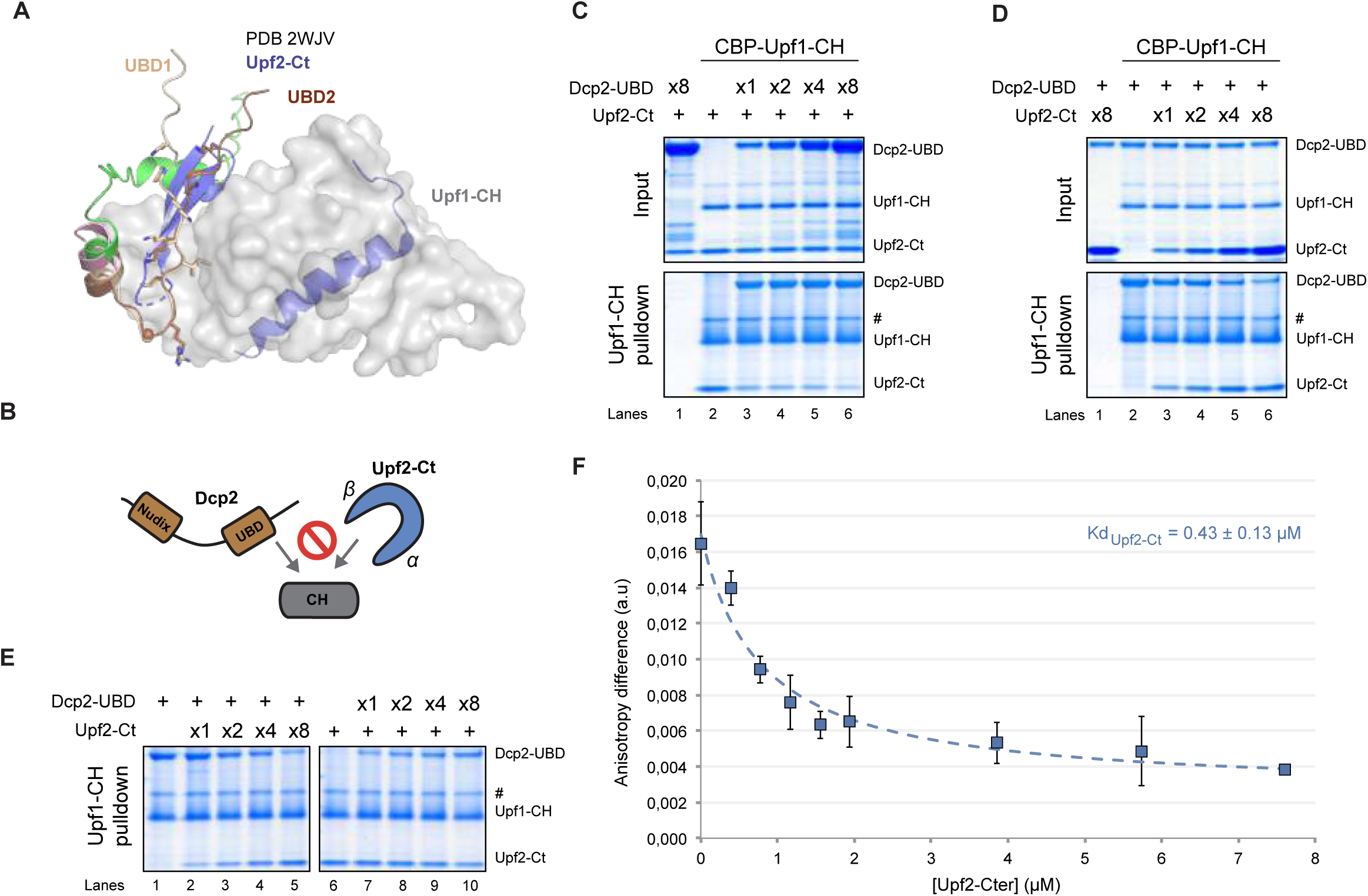
Upf2 and Dcp2 compete for binding to Upf1-CH domain. **(A)** Superposition of the best AF3 models of the *S.c.* Upf1-CH domain (grey, shown as semi-transparent) bound to Dcp2 UBD1 (green) and UBD2 (pink) onto the crystal structure of human Upf1/Upf2 complex ^62^ (PDB code : 2WJV). For the sake of clarity, the human Upf1 CH domain is not shown. The Dcp2 UBD1 (aa 455-475) and UBD2 (aa 694-713) motifs are colored in wheat and brown, respectively. **(B)** Schematic representation of interaction incompatibility between Dcp2 and Upf2 binding to the CH domain of Upf1. **(C, D)** CBP-pulldown assays using Upf1-CH against an increasing amount of Dcp2-UBD-with a constant amount of Upf2-Ct (C) or increasing amount of Upf2-Ct with a constant amount of Dcp2-UBD (D). As previously, protein mixtures before (input, 17% of total) or after precipitation were separated on SDS-PAGE and revealed using Coomassie blue. Protein contaminants are indicated with (#). **(E)** CBP-pulldown assays using pre-formed Upf1-CH/Dcp2-UBD complex against increasing amounts of Upf2-Ct (left) or pre-formed Upf1-CH/Upf2-Ct complex against increasing amounts of Dcp2-UBD (right). **(F)** Chase experiments of FITC-labeled Dcp2-UBD2 peptide (15 nM) bound to CBP-Upf1-CH-His (20 μM) by Upf2-Cter. The curve obtained after fitting of the experimental data with equation (B) from the materials and methods section, is shown as a dashed black line. Error bars were calculated from triplicate experiments.

Consistent with our previous data showing that the Upf1-2/3 and Upf1-decapping complexes purified from yeast are mutually exclusive ^45^, our results suggest that this incompatibility is, at least partly, due to the competition between Upf2 and Dcp2 for interaction with Upf1-CH.

### Upf1 RNA-binding is differentially regulated by its partners

To evaluate the ability of Upf1-FL to bind RNA in the presence or in the absence of its NMD partners, we performed RNA-pulldown assays. As for the protein interaction assays, the proteins were first mixed in the presence of a short (33 mer) 3’-end biotinylated RNA with or without ATP or non-hydrolysable analog ADPNP before purification on streptavidin beads. As previously observed ^16,19,65^, Upf1 binding to RNA differed in the presence or absence of ATP. Upf1-FL was better retained by the RNA in the absence of nucleotide or in the presence of ADPNP, a non-hydrolysable analog ATP, than in the presence of ATP (Figure 5A, lanes 2, 6, 10). Due to unspecific interactions between all Ebs1 tested versions and the streptavidin resin, we did not include this protein in these RNA-pulldown experiments. Moreover, since the TS-tagged version of Nmd4 presented unspecific interactions with the resin, we purified a His-tagged version of Nmd4. In contrast to Nmd4 that was slightly pulled down by RNA, neither Dcp2-UBD nor Upf2-Ct were individually retained by the RNA (Figure 5A, lane 1). The addition of Upf2-Ct reduced by half the amount of Upf1 retained by the RNA (Figure 5A, lanes 5, 9, Figure 5B for quantification). This was particularly visible in the absence of nucleotide or in the presence of ADPNP when Upf1 recovered by RNA purification was the most important, suggesting that Upf2 prevents Upf1 binding to RNA (Figure 5A). Regardless of the condition, Upf2-Ct was never co-precipitated by the RNA in the presence of Upf1 (Figure 5A). Remarkably, we observed an opposite effect when testing two components of the Upf1-decapping complex. The addition of Nmd4 and, to a lesser extent, Dcp2-UBD, significantly increased the amount of Upf1 retained by RNA in the presence of ADPNP or ATP (Figure 5A, lanes 7, 8, 11, 12, Figure 5B). This positive effect was not measurable in the absence of nucleotide as Upf1 was already strongly attached to RNA (Figure 5A, lanes 3, 4, Figure 5B). In contrast to Upf2-Ct, it is noteworthy that Dcp2-UBD and Nmd4 systematically co-precipitated with the RNA-bound Upf1 in all conditions. Therefore, individually, both Dcp2 and Nmd4 stabilize Upf1 bound to RNA. We performed a similar experiment in the presence of ATP by mixing Upf1-FL with either partner, a combination of two partners or all three partners simultaneously (Figure 5C). The most effective retention of Upf1-FL onto RNA was observed in the presence of both Dcp2-UBD and Nmd4, indicating their synergistic contribution to the stability of this complex, with their effects being additive (Figure 5C, lanes 5, 8 and 9, Figure 5D). Upf2-Ct showed no significant effect on the retention of Upf1-FL when either Dcp2-UBD or/and Nmd4 were present (Figures 5C, 5D). These effects were also reproducible in absence of nucleotide (Figure S6A) and in presence of ADPNP (Figure S6B).

**Figure 5:**
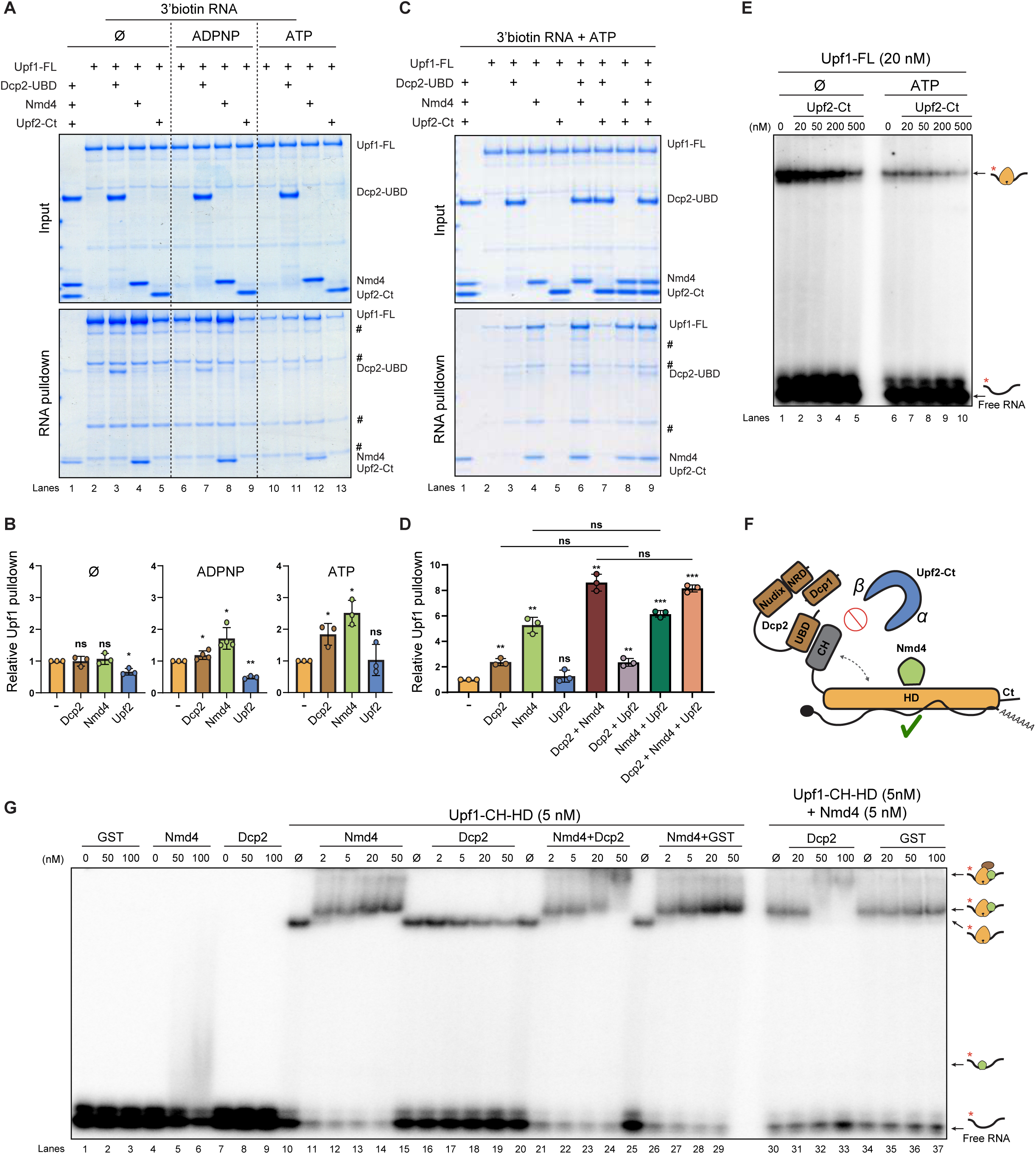
Upf1 RNA-binding activity is modulated by its partners. **(A)** RNA-pulldown assays using a 3’ end-biotinylated 33 mer RNA as bait. As indicated, Upf1-FL, Nmd4, Dcp2-UBD or Upf2-Ct were mixed with the RNA in the absence of nucleotide or in the presence of ATP or ADPNP. Protein mixtures before (input, 17% of total) or after precipitation using magnetic streptavidin beads were separated on SDS-PAGE and revealed using Coomassie blue. Protein contaminants are indicated with (#). **(B)** Relative quantification of Upf1 pulldown in presence of partners (A, lower panel lanes 2-13) normalized by Upf1 pulldown without any partner (A, lower panel lanes 2, 6 or 10). Error bars denote standard deviations between replicates (n=3). Individual measurements are represented as circles, while the mean of each data series is represented as a column. Unpaired *t* tests were performed to calculate *p* values with a significance threshold set at 0.05 marked with one star. *p* values lower than 0.01 were marked with two stars, and lower than 0.001 were marked with three stars. Non-significant p-values were marked with ns. **(C)** RNA-pulldown assays using a 3’ end-biotinylated 33 mer RNA as bait against Upf1-incubated with ATP and combinations of Nmd4, Dcp2-UBD and Upf2-Ct, as indicated. **(D)** Relative quantification of Upf1 pulldown in (C) normalized by Upf1 pulldown without any partner (C, pulldown lane 2). Error bars denote standard deviations between replicates (n=3). Individual measurements are represented as circles, while the mean of each data series is represented as a column. Unpaired *t* tests were performed to calculate *p* values with a significance threshold set at 0.05 marked with one star. *p* values lower than 0.01 were marked with two stars, and lower than 0.001 were marked with three stars. **(E)** Representative native 7% polyacrylamide gel illustrating the interaction of Upf1-FL (depicted in orange) with a ^32^P-labeled 30-mer oligoribonucleotide substrate (black line with red star). The RNA substrate was incubated with 20 nM of Upf1-FL and increasing concentrations of Upf2-Ct and with (+) or without (0) ATP under the conditions described in the materials and methods section. **(F)** Schematic representation of protein-RNA interactions between the Upf1, Dcp1, Dcp2, Nmd4 complex and Upf1-Upf2 complex. **(G)** Representative native 7% polyacrylamide gel illustrating the interaction of Upf1-CH-HD (depicted in orange) with a ^32^P-labeled 30-mer oligoribonucleotide substrate (black line with red star) in presence of Dcp2-UBD (depicted in brown), Nmd4 (depicted in green) and/or GST as a control. The RNA substrate was incubated with 5 nM of Upf1-CH-HD and increasing concentrations of Dcp2-UBD, Nmd4 and/or GST and with ATP under the conditions described in the materials and methods section.

We also performed electrophoretic mobility shift assays (EMSA) to study Upf1 binding to RNA using a complementary approach. We incubated a 30-mer radiolabeled ssRNA with increasing concentrations of CBP-Upf1-CH-HD or CBP-Upf1-FL without nucleotide, ADPNP, or ATP. In each case, a band shift was observed upon gel electrophoresis, indicating Upf1– RNA complex formation (Figure S6C). As previously observed, Upf1 isoforms show more affinity to RNA in the absence of nucleotides ^16,19,65^. We found that both in the absence of nucleotides and in the presence of ATP, increasing amounts of Upf2-Ct do not bind RNA (Figure 5E, lanes 1 and 6), but reduced the Upf1-FL shift by half (Figure 5E, lanes 2-5 and 7-10, Figure S6D), confirming that Upf2 hampers Upf1 binding to RNA in presence or absence of ATP.

Next, we tested the effect of Nmd4 and Dcp2 on Upf1-CH-HD RNA binding. EMSA with Nmd4, Dcp2-UBDs, or a control protein GST added separately, showed only a weak shift when a large excess of Nmd4 (25-fold) was added to RNA (Figure 5G, lanes 4-6). This confirms the weak affinity of Nmd4 for RNA observed by pull-down assays (Figures 5A, 5C). When increasing amounts of Nmd4 were added to a fixed amount of Upf1-CH-HD, Nmd4 produced a super-shift already detected at equimolar concentration with Upf1 (Figure 5G, lanes 11-14). Addition of Dcp2 in the same range showed no super-shift of Upf1 (Figure 5G, lanes 16-20). However, a complete and even higher super-shift was detected when both Nmd4 and Dcp2 were added (Figure 5G, lanes 21-24), while the addition of Nmd4 and GST showed no additional shift (Figure 5G, lanes 26-29). This specific super-shift was confirmed when an equimolar and a fixed amount of Upf1 and Nmd4 were mixed with increasing amounts of Dcp2 but not with the GST control protein (Figure 5G, lanes 30-37).

The EMSA results confirmed our observations from interaction assays (Figures 5A, 5C). Together, these findings demonstrate that Upf1, Nmd4, and Dcp2 form a stable complex on RNA and that Upf1 and Nmd4 are necessary for stably anchoring Dcp2 onto RNA unlike Upf2, which detaches Upf1 from RNA (Figure 5F).

## DISCUSSION

Since its discovery, genetics, functional assays, and biochemical characterization have identified most, if not all, NMD factors in different eukaryotic organisms ^66^. However, the highly dynamic aspect of this process linking premature translation termination with mRNA decay challenges the characterization of the successive molecular steps. Central to NMD, Upf1 recognizes mRNAs containing premature termination codons and coordinates the recruitment of specific decay factors. This involves the formation of transient Upf1-containing complexes that have been only marginally characterized biochemically ^45,51,67,68^. Here, using isolated recombinant yeast proteins we could recapitulate critical phases of NMD and notably, direct interaction and competition between NMD and decay factors and their binding to RNA.

All three distinct domains of Upf1, the N-terminal CH domain, the central helicase domain (HD), and the disordered C-terminal domain, make direct contacts with Dcp2, Dcp1, Nmd4, and Ebs1 (Figures 1, S1, S3). Interestingly, many of the partners we included in our study directly contact distinct domains of Upf1 and other additional secondary interactions between other partners take place within the complex (Figure 2). These findings rationalize the fact that deletion of single contact points or partners is insufficient to prevent the assembly of the complex and accounts for the weak NMD target destabilization upon individual Ebs1 and Nmd4 deletions or deletion of the C-terminal domain of Dcp2 ^45,47^. The CH domain of Upf1 holds a pivotal position in this intricate network of interactions as it contacts all tested partners, notably the decapping protein Dcp2 and Upf1’s essential co-factor Upf2. Dcp2 directly interacts with Upf1 through two Upf1-binding domains, UBD1 and UBD2 ^47,51,52^. We show that both UBD domains contact the same hydrophobic pocket within Upf1 CH domain and that UBD2 possesses a higher affinity than UBD1 from binding Upf1-CH domain (Figure 1). Since deletion of the UBD domains scarcely impacts NMD efficacy *in vivo* ^47^, it remains unclear whether both UBDs contribute to the architecture of Upf1-containing decapping complexes. However, the lack of impact of UBD presence on the function of the Upf1-decapping complex can be explained by taking into consideration the previously unknown role of Dcp1, which binds directly to Upf1, as shown here (Figure 1E).

Our biochemical approach revealed that the interactions of Dcp2 UBDs with the Upf1 CH domain are mutually exclusive with Upf2 binding (Figure 4). Dcp2 UBDs are predicted by AF3 to contact the same external side of the CH domain that binds the β hairpin of Upf2 (Figure 1 and S1). Two conserved regions in Upf2, a β hairpin and an α helix, clamp two opposite sides of the Upf1 CH domain ^62^ (Figure 3). This bipartite mode of interaction is therefore conserved between yeast and human ^62,69^. Interaction assays using targeted mutagenesis further confirmed this model (Figure 3). The hydrophobic pocket of the CH domain is involved in the interaction with the Upf2 α-helix and contacts conserved residues of the RecA2 domain and the stalk—two structural elements of the helicase domain. The Upf1 I693R mutation on the surface of RecA2 favors the interaction of Upf2 most likely by giving access to both binding sites. Conversely, the F131E mutation in the hydrophobic pocket of the CH domain strongly reduces Upf2 interaction indicating the residue is involved in the interaction (Figure 3). In human Upf1, the corresponding residue is also implicated in the interaction with Upf2 ^16^. The biochemical evidence for the mutually exclusive binding of Upf1 to Upf2 or Dcp2 accounts for the two distinct Upf1-2/3 and Upf1-decapping NMD complexes isolated *in vivo* ^45^. This transition between the Upf complex and the Upf1-decapping complex, which triggers mRNA decay orchestrated by the CH domain of Upf1, is most likely also conserved. In yeast, NMD targets are mainly degraded by decapping ^41^, making Dcp2 the main decay-inducing partner of Upf1. In human, the predominant RNA degradation pathway in NMD involves the endonucleolytic cleavage of mRNA by Smg6 ^36–38,70^. Interestingly, recent biochemical evidences revealed that Smg6 and Upf2 also compete for their binding to the CH domain of Upf1 in human ^71^ supporting a sequential binding of these factors during NMD similar to Dcp2 and Upf2 in yeast.

Our biochemically reconstituted system allowed determining how Upf1 direct partners influence Upf1 binding to RNA (Figure 5). RNA-based pulldowns and EMSA experiments first showed that Upf2 reduces Upf1 binding to RNA. Yeast Upf1 HD domain contacts RNA in a closed Upf1 conformation that is controlled by the relative orientation of the CH and HD domains ^16^. Docking the Upf1 CH domain onto the HD domain increases the surface contacted by RNA. This docking cannot occur when opposite faces of the CH domain are clamped by Upf2. This negative impact of Upf2 on Upf1 affinity to RNA is conserved as a similar effect occurs with the human proteins ^58,72^. Given that Upf2 bridges Upf1 and Upf3 ^58^, it is tempting to speculate that as part of the Upf1-2/3 complex, Upf1 is not stably attached to the RNA. Conversely, Nmd4 stabilizes Upf1 bound to RNA allowing in turn Upf1 to tether Dcp2 to the RNA. In contrast to Upf2, Dcp2 only binds the external face of the CH domain and thus does not prevent its docking to the HD domain in the closed conformation of Upf1. Dcp2 and Nmd4 likely promote Upf1 RNA-binding by stabilizing this conformation. Therefore, we provide evidence of two distinct states of Upf1 regarding its binding to RNA during the stepwise NMD pathway: a poor affinity to RNA in the presence of Upf2 and its stable binding to RNA when bound to the components of the NMD-decapping complex. If the negative effect of Upf2 on Upf1 binding to RNA is conserved in human, it is tempting to believe that Upf1 also contributes to stabilize the association of the decay factors with the RNA target during the late phases of NMD.

These biochemical snapshots of Upf1-containing complexes offer a more precise view of the cascade of Upf1-dependent events leading to PTC-containing mRNA decay (Figure 6). The delayed process of release factors recruitment and ribosome recycling upon premature translation termination, favor the recruitment of Upf1 to the terminating ribosome ^73–75^. This contact is facilitated by the promiscuous Upf1 binding to RNA enriched onto mRNA 3’UTR ^2,76–78^. This step coincides with Upf1-dependent recruitment of the Upf2/Upf3 heterodimer ^45,79^ that also contributes to delay translation termination ^80^. In the Upf1-2/3 complex, Upf1 would no longer be strongly bound to RNA ^58,72^. Next, a molecular switch that remains to be characterized displaces Upf2/3 thus allowing assembly of the Upf1-decapping complex ^45^. Contact with the nearby RNA and assembly of the decapping complex may contribute to detaching Upf2 from Upf1, as Upf1’s interaction with Upf2 is incompatible with RNA and Dcp2 binding (Figure 5). In the Upf1-decapping complex, Upf1 stably bound to RNA in the presence of its co-factors, anchors the major decay factors Dcp1-Dcp2 establishing the irreversible step towards RNA degradation.

**Figure 6:**
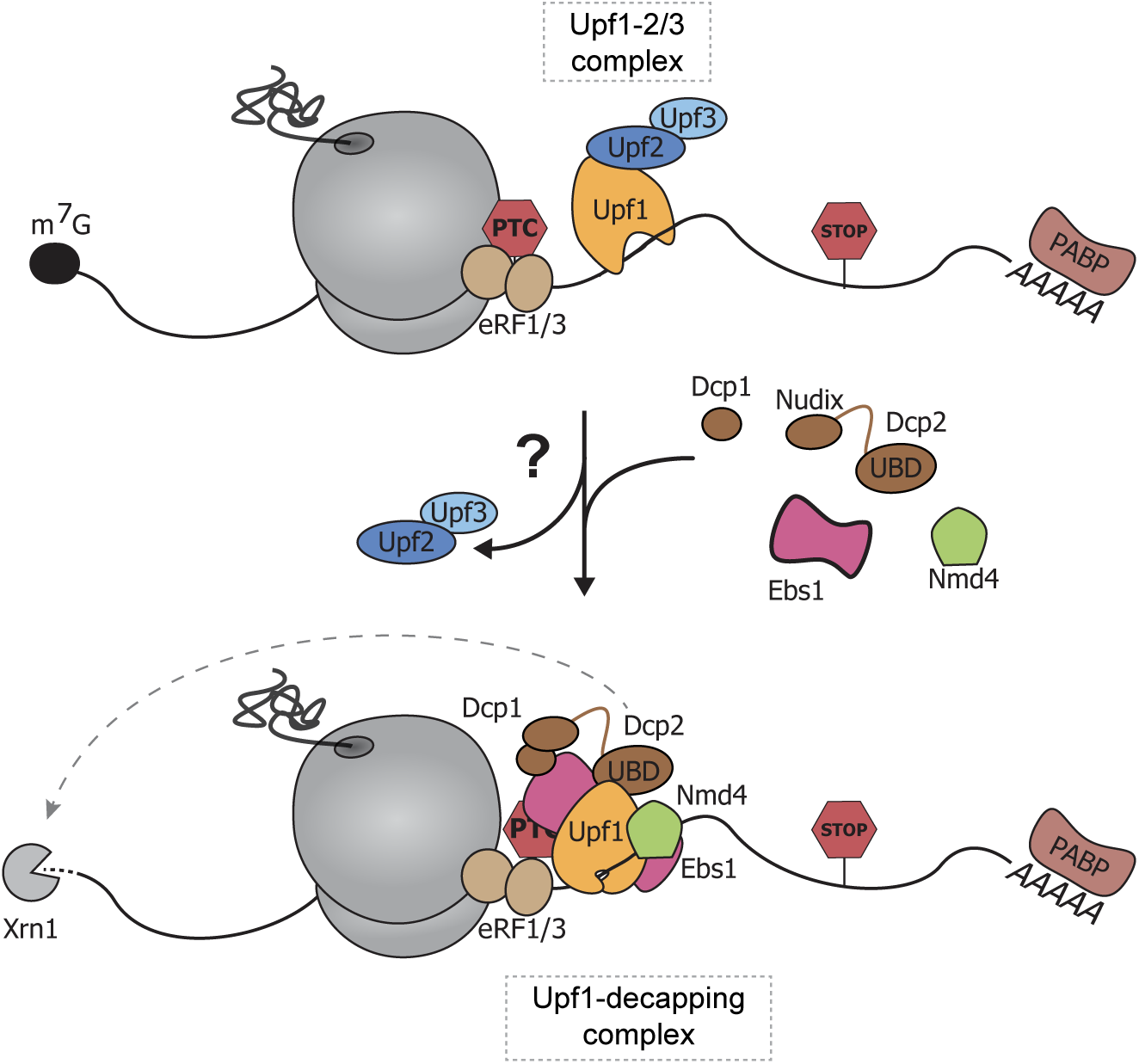
Proposed model for Upf1-2/3 and Upf1-decapping complex sequential formation during NMD in yeast

The *in vitro* approaches employed in this study allowed zooming in on important molecular interactions between Upf1, its NMD partners, the decay machinery and the RNA. These insights pave the way for more extensive structural studies and improve our understanding of conserved NMD mechanisms in eukaryotes. However, this experimental strategy faces several limitations such as the difficulty to produce high yields of full-length isoforms for each protein and thus to include all NMD factors, such as Upf3. So, if our strategy allows describing important interactions, we cannot exclude complementary or alternative interactions between NMD factors. Although we expressed multiple recombinant protein isoforms, we may have missed additional contacts offered by full-length and post-translationally modified proteins. Other factors that we did not include in our reconstitution may have important roles, such as the decapping co-factor Edc3, which facilitates the association of Upf1 with Dcp2 *in vivo* and activates Dcp2 ^45,47,50,81^, or the protein kinase Hrr25, which likely transiently interacts with Upf1 and may participate in post-translational modifications ^45^. Moreover, NMD is a highly dynamic process and transitions between successive complexes from translation termination and RNA decay as well as the modulation of enzymatic activities are challenging to catch *in vitro*. For example, the molecular determinant operating the switch between the Upf1-2/3 detector complex and the Upf1-decapping complex will be key to understand the NMD pathway.

## Materials and methods

### cDNA cloning

*Saccharomyces cerevisiae* (*S.c.*) Upf1 full length (FL, pHL1280), helicase domain (HD, pHL1281) and CH-HD truncation (pHL 1282) (Uniprot accession code P30771) were cloned into a homemade variant of pET28a vector (Novagen) containing Calmodulin-Binding Protein (CBP) coding sequence (pHL5) and hexa-histidine tag (6-His or His) coding sequence. PCR products were amplified using HLH2703/HLH2704, HLH2705/HLH2706 and HLH2707/HLH2706 respectively and were inserted between NheI/XhoI. Truncations (CH, HD-Ct and Ct, respectively pHL1603, pHL1559 and pHL1566) were cloned by PCR amplification using phosphorylated oligonucleotides followed by ligation using HLH3765/HLH2824, HLH3641/HLH3642 and HLH3641/HLH3643 as oligonucleotides respectively and pHL1280 as matrix. Domain boundaries were defined according to previous studies ^16^.

S.c. Dcp2-UBD (pHL1637) (Uniprot accession code P53550) was amplified using HLH3826/HLH3841 and cloned into pGEX-6P-1 vector (Cytiva) between BamHI/NotI. *S.c.* S.c. Dcp2-UBD1 (aa 434-508, pMG621) and UBD2 (aa 690-720, pMG676) were also cloned into pGEX-6P-1 vector between BamHI/NotI. GST-Dcp1 (pBS4576) and GST-Dcp1/Dcp2-S aa 1-315, pBS5135) were generated as described in ^50^. Dcp2(1-663) and Dcp2(1-720) were cloned into pBS5135.

*S.c.* Nmd4 full-length coding sequence (Uniprot accession code Q12129) was provided by C. Saveanu ^45^. The sequence was cloned into a homemade variant of pET28a vector (Novagen) containing Twin-Strep-tag (TS) coding sequence (pHL1567) to produce TS-Nmd4-His. It was cloned into pHL5 to produce CBP-Nmd4-His. To produce the C-terminal His-tagged version, it was cloned into pet28a. Finally, the pMG838 plasmid expressing the HisZZ-tagged Nmd4 protein was generated as described in ^82^.

S.c. Ebs1(1-591) coding sequence (Uniprot accession code Q03466) was provided by C. Saveanu ^45^ and cloned into a homemade variant of pET28a (Novagen) containing Glutathion-S-Transferase coding sequence (pHL6) to obtain pHL1513 or CBP (pHL5) to obtain pHL1514. Both PCR products were amplified using Ebsrev/Ebsfw and inserted between NdeI and XhoI. To produce the His-SUMO-tagged version, Ebs1(1-591) was amplified using HLH3801/HLH3802 and inserted between NotI and XhoI into a modified pBR322 vector with His-SUMO tag at the 3’ end ^83^.

S.c. Upf2(933-1089) coding sequence (Uniprot accession code P38798) was amplified using Upf2-933-NdeI-for/Upf2-1089-NotI-rev and cloned between NdeI/NotI into a modified pET9b vector with an addition of a sequence coding for an hexahistine tag at the 3’ end of the gene (pMG462).

The CBP-HisZZ plasmid was cloned by Gibson assembly using a gene block containing CBP coding sequence ordered from IDT inserted between NcoI and NdeI within a pet28b vector containing the HisZZ coding sequence ^82^.

Oligonucleotides:

HLH2703 (ctagacgctagcatggtcggttccggttctcacac) HLH2704 (catgacctcgagtattcccaaattgctgaagtcttttgac) HLH2705 (ctagacgctagcgaacaggaagcaatcccacc) HLH2706 (catgacctcgagaggacgaactaattgaacagtgc) HLH2707 (ctagacgctagctcgccttcagcttcagac) HLH3765 (ctctggggcgtcaatatcattaatt)

HLH2824 (ctcgagcaccaccaccacc) HLH3641 (gctagccatatgctccttatcg) HLH3642 (gaacaggaagcaatcccacc) HLH3643 (cagccaagaaagactgaacg) HLH3826 (cctgggatccagcagctcctcccctgg)

HLH3841 (cgatgcggccgcaagcttttttatcattcgccagac)

Upf2-933-NdeI-for (gggaacatatgagcgactctgatttggagtatggtgg)

Upf2-1089-NotI-rev (gggaacatatgcaccatcaccatcaccatgttcgaaagaacgttttaaaacaatctttttaatcttgtttcgc) Ebsfw (cccatatggctagcatggaaccatcga)

Ebsrev (gtggtgctcgagcttaaatccaa)

HLH3801 (aaaccggcggccgcatggaaccatcgaatacccaaaaag) HLH3802 (tcacgttatatctttaataagaaaattggatttaagtgactcgagcaccac)

### Protein overexpression and purification

BL21(DE3)-CodonPlus competent *Escherichia coli* bacteria (Agilent) were transformed with 500 ng of plasmid of interest by 2 min at 42°C heat shock. After addition of 1 mL liquid Lysogeny Broth (LB) media and 1 hour incubation at 37°C, cells were pelleted by 4 min centrifugation at 3500 g. The pellet was resuspended in 50 µL LB media, plated in antibiotic containing LB plates and left to grow overnight at 37°C. 2-3 colonies were resuspended in 25 mL LB liquid media containing antibiotic in a 250 mL Erlenmeyer flask and incubated at 37°C, 200 rpm for 4 hours. The bacteria were inoculated in 1 L of autoinducible LB or TB (Terrific Broth) containing metals (Formedium) added with the plasmid antibiotic resistance (ampicillin 100 µg/mL or kanamycine 50 µg/mL) and chloramphenicol (25 µg/mL) in a 5 L Erlenmeyer flask and incubated at 18°C, 200 rpm overnight. Bacterial cultures were pelleted by 10 min, 4 000 g centrifugation at 4°C. The pellet was resuspended in 25 mL of His-Lysis Buffer (1.59 mM KH_2_PO_4_, 4.5 mM, Na_2_HPO_4_, 231 mM NaCl, 1 mM MgAc2, 0.1% NP-40, 20 mM imidazole, 10% (v/v) glycerol, 2 mM DTT, pH 7.0). Before sonication, 1X of protease inhibitors (Apoprotinine, Leupeptine, PMSF, Pepstatin) were added. Samples were sonicated in a metal beaker at 30% amplitude and 1 second pulse during 4 min. In order to keep low temperatures the beaker was placed on a mix of dry ice and ice. The bacterial debris were pelleted by centrifugation for 30 min at 18 000 rpm. The supernatant was retrieved and mixed with 1 mL of Nickel-NTA Agarose slurry (Qiagen) equilibrated with His-Lysis Buffer. The sample was incubated in a rotator for 1 hour at 4°C before centrifugation at 4 000 rpm for 5 min at 4°C to pellet the beads. The supernatant was removed and the beads were washed with 25 mL of His-Lysis Buffer. The beads were transfered into a Poly-Prep chromatography column (Bio-Rad) pre-equilibrated with 1 mL of Lysis Buffer. The column was washed with 5 mL of His-Lysis Buffer then with 5 mL of His-Wash Buffer (1.59 mM KH_2_PO_4_, 4.5 mM, Na_2_HPO_4_, 231 mM NaCl, 250 mM NaCl, 1 mM MgAc_2_, 0.1% NP-40, 50 mM imidazole, 10% (v/v) glycerol, 2 mM DTT, pH 7.0) then again with 5 mL of His-Lysis Buffer. The column was closed and the resin was incubated with 1 mL of His-Elution Buffer (1.59 mM KH_2_PO_4_, 4.5 mM, Na_2_HPO_4_, 231 mM NaCl, 1 mM MgAc_2_, 0.1% NP-40, 250 mM imidazole, 10% (v/v) glycerol, 2 mM DTT, pH 7.0) for 10 minutes at 4°C. Elution was performed successively with 1 mL of Elution Buffer and protein concentrations were measured by Bradford Assay. The fractions with highest concentration were pooled together and dialysed overnight. For CBP-tagged proteins, dialysis was performed against 1 L of Calmodulin Binding Buffer (CBB) (10 mM Tris-HCl pH 7.5, 250 mM NaCl, 1 mM MgAc_2_, 4 mM CaCl2, 0.05% NP-40, 10% (v/v) glycerol, 2 mM DTT) at 4°C. The dialysed fractions were retrieved and mixed with 500 µL of Calmodulin Sepharose 4B resin (50% Slurry, GE Healthcare) washed with CBB. The samples were incubated in a rotor for 2 hours at 4°C then transferred into a Poly-Prep chromatography column (Bio-Rad) pre-equilibrated with 1 mL of CBB. After one wash with 5 mL of CBB the column was closed and the resin was incubated with 500 µL of Calmodulin Elution Buffer (10 mM Tris-HCl pH 7.5, 250 mM NaCl, 1 mM MgAc_2_, 0.05% NP-40, 20 mM EGTA, 10% (v/v) glycerol, 2 mM DTT) for 10 min at 4°C. Elution was performed successively with 500 µL of Calmodulin Elution Buffer and protein concentrations were measured by Bradford Assay. For GST- and TS-tagged proteins, purification steps were similar but performed using different buffers. For GST-tagged proteins, dialysis or lysis was performed against GST-Binding Buffer (20 mM Tris-HCl pH 8.0, 200 mM NaCl, 5mM β-mercaptoethanol, 1 mM MgCl_2_), the resin used was Glutathione Sepharose 4B (GE Healthcare), and proteins were eluted with GST Elution Buffer (50 mM Tris-HCl pH 7.5, 250 mM KCl, 1 mM MgAc2, 1 mM DTT, 0.005% NP-40, 10% (v/v) glycerol, 10 mM L-glutathione reduced). For TS-tagged proteins, dialysis was performed against TS-Binding Buffer (10 mM Tris-HCl pH 8, 150 mM NaCl, 0.1% NP-40, 1 mM DTT). StrepTactin resin (Iba) was used for protein binding and proteins were eluted with TS Elution Buffer (10 mM Tris-HCl pH 8, 150 mM NaCl, 0.1% NP-40, 1 mM DTT, 2.5 mM D-Desthiobiotin). For all eluted proteins the highest concentrated fractions were pooled together and dialysed overnight against 1 L of Storage Buffer (1.59 mM KH_2_PO_4_, 4.5 mM, Na_2_HPO_4_, 231 mM NaCl, 1 mM MgAc2, 10% (v/v) glycerol, 2 mM DTT, pH 7.0). The dialysed proteins were aliquoted in small volumes depending on the final protein concentration and stored at -80°C ^84,85^.

The S.c. Upf2(933-1089)-His protein was over-expressed from pMG462 in *E. coli* BL21 (DE3) CodonPlus cells in 1 L of TBAI containing kanamycin (100 µg/mL) and chloramphenicol (25 µg/mL). After a 3 h incubation at 37 °C, the bacteria were incubated overnight at 18°C. Cells were harvested by centrifugation at 4,000 rcf for 30 minutes at 4°C, resuspended in buffer Lysis Buffer B (20 mM Tris-HCl pH 8.0, 200 mM NaCl, 5 mM β-mercaptoethanol) and lysed by sonication after addition of 200 µL of phenylmethylsulfonyl fluoride (PMSF; 100 mM). The protein was first purified on a Protino® Ni-NTA agarose resin pre-equilibrated with buffer Lysis Buffer B. After successive washing steps using first Lysis buffer B and then « high-salt » buffer (20 mM Tris-HCl pH 8.0, 1 M NaCl, 5 mM β-mercaptoethanol), the proteins were eluted using Lysis buffer B supplemented with 400 mM imidazole pH 7. The eluted protein was loaded on a HiTrap Heparin (Cytiva) column pre-equilibrated in buffer A_50_ and eluted using a linear gradient from 100 % of « low-salt » buffer (20 mM Tris-HCl pH 8.0, 50 mM NaCl, 5 mM β-mercaptoethanol) to 100 % of « high-salt » buffer. Finally, the protein was purified on a S75-16/60 size-exclusion column (Cytiva) equilibrated with Lysis buffer B supplemented with 10% (v/v) glycerol.

### The HisZZ-tagged Nmd4 protein was purified as described in ^82^

The CBP-His-ZZ protein was over-expressed from pHL1662 in *E. coli* BL21 (DE3) Gold cells in 1 L of TBAI containing kanamycin (100 µg/mL). After a 3 h incubation at 37 °C, the bacteria were incubated overnight at 25°C. Cells were harvested by centrifugation at 4,000 rcf for 30 minutes at 4°C, resuspended in buffer Lysis Buffer C (20 mM Tris-HCl pH 7.5, 200 mM NaCl) and lysed by sonication after addition of 200 µL of phenylmethylsulfonyl fluoride (PMSF; 100 mM). The protein was first purified on a Protino® Ni-NTA agarose resin pre-equilibrated with buffer Lysis Buffer C. After successive washing steps using first Lysis buffer C and then high-salt buffer (20 mM Tris-HCl pH 7.5, 1 M NaCl), the proteins were eluted using Lysis buffer C supplemented with 250 mM imidazole pH 7. The eluted protein was loaded on a HiTrap Q (Cytiva) column pre-equilibrated in Lysis buffer C and eluted using a linear gradient from 100 % of Lysis buffer C to 100 % of high-salt buffer. Finally, the protein was purified on a S75-16/60 size-exclusion column (Cytiva) equilibrated with Lysis buffer C.

### Protein and RNA pulldown assays

6μg of each protein of interest were mixed together in a Protein LoBind tube (Eppendorf) and volume was completed to 30 μL with Storage Buffer (1.59 mM KH_2_PO_4_, 4.5 mM, Na_2_HPO_4_, 231 mM NaCl, 1 mM MgAc2, 10% (v/v) glycerol, 2 mM DTT, pH 7.0). A 5 μL sample was retrieved from each tube as input and mixed with 10 μL of protein loading dye (LD) 4X. For RNA free pulldowns, 5 μL of H_2_O were added. For RNA-pulldown assays, 1 μL of H_2_O, 2 μL of 10 mM 3’ biotinylated 30 mer RNA (rCrGrU rCrCrA rUrCrU rGrGrU rCrArU rCrUrA rGrUrG rArUrA rUrCrA rUrCrG, ordered from IDT) and 2 μL of 50 μM ADPNP or ATP were added. The samples were complemented with 30 μL of Binding Buffer 2X (40 mM Hepes pH 7.5, 87 mM NaCl, 4 mM MgAc2, 4 mM Imidazole, 0.2% NP-40, 4 mM DTT, and 4 mM CaCl2 if CBP pull-down) and incubated for 20 minutes at 30°C. For CBP pulldowns assays, 15 μL of Calmodulin Sepharose 4B resin (50% Slurry, GE Healthcare) were added. For RNA-pulldowns, 5 μL of preblocked Streptavidin Dynabeads MyOne (Thermofisher) were added. Beads were pre-blocked with 40 μL of 5 M NaCl, 2.5 μL of Glycogen Carrier (2 mg/mL), 5 μL of tRNA yeast (10 mg/mL), 50 μL of BSA (10 mg/mL). The samples were completed with Binding Buffer 1X (20 mM Hepes pH 7.5, 125 mM NaCl, 2 mM MgAc2, 2 mM Imidazole, 0.1% NP-40, 2 mM DTT, and 2 mM CaCl2 if CBP pulldown) to make a total volume of 200 μL and incubated for 1 hour at 4°C with rotation. The beads were washed three times with 800 μL of Wash Buffer 1X (20 mM Hepes pH 7.5, 2 mM MgAc2, 2 mM Imidazole, 0.1% NP-40, 2 mM DTT, and 2 mM CaCl2 if CBP pulldown, complemented with either 250 mM, 300 mM, 350 mM or 500 mM NaCl as mentioned). After each wash, samples were rotated for 2 minutes at 4°C, centrifuged for 30 seconds at 2 000 rpm and 4°C. Proteins were eluted with 10 μL of LD 4X and loaded into a 15-well NuPAGE 4-12% acrylamide SDS-PAGE (Thermo Fisher Scientific). Gels were scanned using an Eson Perfection V700 Photo scanner and quantification of protein pulldown was performed using FIJI Gel Analyzer tool to measure the signal value of pulldown gels. Values were normalized as mentioned in figure legends and unilateral unpaired Student t tests were performed to assess statistical significance.

### AlphaFold3 simulations

Five 3D structure models of S.c. Upf1-CH bound to Dcp2-UBD1, Dcp2-UBD2 or Upf2-Ct were generated using the seed auto option of the recently launched AlphaFold3 server (AF3; version 3.0; https://alphafoldserver.com; ^63^) and the aa sequences of the proteins of interest ^63,86^. The 3D model of the S.c. Ebs1 protein was generated using the same approach.

### Sequence alignments

Multiple sequence alignment of fungal Dcp2 proteins was obtained by performing PSI-BLAST using the sequence of *S.c.* Dcp2 ^87^. The identification of the UBD1 and UBD2 motifs was made easier thanks to former yeast two-hybrid data that have identified conserved residues in two Dcp2 regions interacting with Upf1 ^51^.

### Electrophoretic mobility shift assay (EMSA)

40 pmoles of a single-stranded 30-mer oligoribonucleotide ^88^ was 5’end-labeled using 80 U of T4 polynucleotide kinase (New England Biolabs) and 10 fmoles of □32P-ATP (3000 Ci/mmole, Perkin Elmer). EMSA samples were prepared by mixing 4 nM of the radiolabelled RNA with the indicated concentrations of proteins, with or without ADPNP or ATP (1 mM) in a buffer containing 20mM MES pH 6.0, 100mM potassium acetate, 2mM DTT, 0.2 mg/ml of BSA and 6% (v/v) glycerol. The 10 µl reaction mixtures were incubated at 30**°**C for 20 min before incubation on ice and addition of 1 µl of loading dye (0.05% xylene cyanol, 0.05% bromophenol blue and 10% (v/v) glycerol). Half of the samples were loaded on a 0.8 mm thick native 7% polyacrylamide (29:1) gel containing 0.5% TBE. Electrophoresis was run for 2.5 hours, constant voltage (240 volts) before exposure of the gel at -50°C and analysis by phosphorimaging (Typhoon FLA 9000 gel imager, GE).

### Fluorescence anisotropy

N-terminally FITC-labeled Dcp-UBD1 (aa 456-479) and Dcp-UBD2 (aa 694-718) peptides were purchased from Proteogenix. Quality and purity of the final products were assessed by HPLC and mass spectrometry.

Fluorescence anisotropy measurements were performed using a FP8300 spectrofluorimeter (Jasco) at 20°C. Excitation and emission wavelengths were 495 nm and 520 nm, respectively. Excitation and emission slits were set-up at 2.5 nm and 5 nm, respectively. Response time was set to 1 second. Each anisotropy value corresponds to the mean value obtained from 15 measurements.

Binding curves were determined by titrating the fluorescent peptides with increasing amounts of CBP-Upf1-CH-His (0 to 40 μM) or CBP-His-ZZ (0 to 35 μM) into a solution composed of 20 mM HEPES pH 7.5, 125 mM NaCl and 2 mM MgAcetate and containing the respective fluorescent Dcp2-UBD peptide (UBD) at a concentration of 15 nM in a 700 μL final volume. Kd_UBD_ and dR_max_ (maximum value of anisotropy difference) were calculated by fitting experimental curves with equation (A) using the program OriginLabs:

**(A)** dR=(dR_max_* [Upf1-CH])/(Kd_UBD_ + [Upf1-CH])

where dR is the anisotropy difference for a given Upf1-CH concentration ([Upf1-CH]).

For competition experiments, the fluorescent Dcp2-UBD2 peptide (15 nM) was first incubated with CBP-Upf1-CH-His (20 μM) and then titrated with increasing concentrations of Upf2-Cter. Dissociation constant (Kd_Upf2-Cter_) was determined by fitting the experimental curve with equation (B):

**(B)** dR=((dR_max_-dR_min_) * [Upf1-CH])/([Upf1-CH] + Kd_UBD2_(([Upf2-Cter]/Kd_Upf2-Cter_) +1)))+dR_min_ where dR is the anisotropy difference for a given concentration of unlabeled Upf2-Cter ([Upf2-Cter]).

**Figure Supplementary 1**

**(A, B, C, D, E)** CBP-pulldown assays using CBP-tagged Upf1 truncations against GST as a control of unspecific interaction (A), Dcp1/Dcp2-L (B), Dcp1/Dcp2-S (C), Dcp2-UBD1 (D), or Dcp2-UBD2 (E). Protein mixtures before (input, 17% of total) or after precipitation (pulldown) were separated on SDS-PAGE and revealed using Coomassie blue. The CH dimer is indicated with (*) and protein contaminants with (#).

**(F)** CBP-pulldown competition assays using Upf1-CH as bait against Dcp2-UBD1 in excess of GST (left) and Dcp2-UBD2 in excess of GST (right). The excess in input of GST is marked in the top panel.

**(G)** Quantification of relative band intensities for Dcp2-UBD1 (wheat) and Dcp2-UBD2 (brown) pull-downs in the presence of increasing amounts of competitor (x1, x2, x4, x8). Mean values are represented as circles, with error bars indicating the standard deviation (n=3). Data were normalized to the signal intensity of pull-down without competitor (Figure 1G, pulldown lanes 2 and 8).

**(H)** Quantification of relative band intensities for Dcp2-UBD1 (wheat) and Dcp2-UBD2 (brown) pull-downs in the presence of x1 competitor with varying excess amounts (x1, x2, x4, x8). Mean values are represented as circles, with error bars indicating the standard deviation (n=3). Data were normalized to the signal intensity of pull-down with x1 excess (Figure 1G, pulldown lanes 3 and 9).

**(I)** Kd values for FITC-labeled Dcp2-UBD1 (wheat) or Dcp2-UBD2 (brown) peptides were determined by fluorescence anisotropy by titration with CBP-Upf1-CH-His (filled squares; n=3). Control experiments were performed by titrating each FITC-labeled Dcp2-UBD peptide with increasing concentrations of CBP-HisZZ (filled triangles; n=2).

**Figure Supplementary 2**

**(A, B)** Multiple sequence alignments of Dcp2-UBD1 (A) and Dcp2-UBD2 (B) domains in the S*accharomycetaceae* family. Strictly conserved residues are in white on a red background. Partially conserved amino acids are shown in red. The consensus motifs are shown below the alignment. This figure was generated using the ESPript server (https://espript.ibcp.fr) ^90^.

**(C, D)** Overall view of the AF3 models generated for the Upf1-CH domain (shown as grey surface) bound to Dcp2-UBD1 (C) and Dcp2-UBD2 (D). Each of the five models is colored differently as indicated. The Dcp2 UBD1 (aa 455-475) motif is colored in wheat and Dcp2-UBD2 (aa 694-713) in brown. The side chain of conserved residues between UBD1 and UBD2 are shown as sticks. The C⍺ atom of the conserved glycin is shown as a sphere.

**(E, F)** Predicted aligned error (PAE) plot of the model of Upf1-CH/Dcp2-UBD1 AF3 model (E) and Upf1-CH/Dcp2-UBD2 model (F).

**Figure Supplementary 3**

**(A)** Cartoon representation of the AlphaFold3 model of *S. cerevisiae* Ebs1 colored by pLDDT values (up) and 14.3.3 domain (aa 1-591) colored in purple and the C-terminal extension is shown in grey (down).

**(B, C)** CBP-pulldown assays using Upf1 truncations against Nmd4 (B) or Ebs1 (C). Protein mixtures before (input, 17% of total) or after precipitation (pulldown) were separated on SDS-PAGE and revealed using Coomassie blue. CH dimer is indicated with (*) and protein contaminants with (#).

**(C)** GST-pulldown assays using Dcp1-FL/Dcp2-L as bait against combination of Nmd4, Ebs1 and Upf1-FL as indicated.

**(B)** CBP-pulldown assay using CBP-Ebs1 as bait against Dcp2-UBD, Dcp1-FL/Dcp2-L2 or isolated Dcp1-FL.

**(C)** TS-pulldown assay using TS-Nmd4 as bait against Dcp1-FL/Dcp2-L2 or isolated Dcp1-FL.

**Figure Supplementary 4**

**(A)** Overall view of the AF3 models generated for the Upf1-CH domain (shown as grey surface) bound to Upf2-Ct. Each of the five models is colored differently as indicated.

**(B)** Predicted aligned error (PAE) plot of the model of Upf1-CH/Upf2-Ct.

**Figure Supplementary 5**

**(A)** Relative band intensity quantification of Upf2 (blue) and Dcp2 (brown) pulldown (Figures 4C and 4D) in presence of increasing quantities of competitor (x1, x2, x4, x8 on the x-axis) normalized to the signal measured in absence of competitor (Figure 4C, pulldown lane 2 and Figure 4D pulldown lane 2). Mean measurements are represented as circles and error bars correspond to standard deviation of replicates (n=3).

**(B)** Relative band intensity quantification of Upf2 (blue) and Dcp2 (brown) pulldown (Figures 4C and 4D) in varying excess amounts (x1, x2, x4, x8 on the x-axis) in presence of x1 of competitor normalized to the signal measured in 1x excess (Figure 4C, pulldown lane 3 and Figure 4D pulldown lane 3). Mean measurements are represented as circles and error bars correspond to standard deviation of replicates (n=3).

**Figure supplementary 6**

**(A, B)** RNA-pulldown assays using a 3’ end-biotinylated 33 mer RNA as bait. As indicated, Upf1-FL, Nmd4, Dcp2-UBD or Upf2-Ct were mixed with the RNA in the absence of nucleotide or in the presence of ADPNP (A) or in absence of nucleotide (B). Protein mixtures before (input, 17% of total) or after precipitation using magnetic streptavidin beads were separated on SDS-PAGE and revealed using Coomassie blue. Protein contaminants are indicated with (#). Relative Upf1 pulldown signal quantification (lower panel) normalized to lanes 4 of both experiments are shown.

**(C)** Representative native 7% polyacrylamide gel illustrating the interaction of Upf1-CH-HD and Upf1-FL (depicted in orange) with a ^32^P-labeled 30-mer oligoribonucleotide substrate (black line with red star). The RNA substrate was incubated with increasing quantities of Upf1 isoforms with (+) or without (0) ATP or ADPNP under the conditions described in the materials and methods section.

**(D)** Relative quantification of Upf1-RNA shift (Figure 5E) in absence (blue circles, straight line) or presence of nucleotide (blue squares, dotted line) normalized by Upf1-RNA shift in absence of Upf2-Ct (Figure 5E lanes 1 and 6 respectively). Mean measurements are represented as circles and error bars correspond to standard deviation of replicates (n=3).

Upf1 is recruited to the target mRNA vicinity adjacent to the terminating ribosome, facilitated either through self-association or interaction with the ribosome within the Upf1-2/3 complex. A pivotal conformational shift ensues, facilitating Upf1 RNA-binding and the assembly of the Upf1-decapping complex, in which Nmd4 which strongly stabilizes Upf1 and Dcp1/Dcp2 onto the mRNA and to initiate mRNA degradation.

## Acknowledgements

We thank Z. Fourati, and B. Seraphin for plasmids. F. Bonneau, G. Badis-Breard, F. Fiorini and members of our laboratory for insightful discussions. O. Bensaude and B. Seraphin for manuscript proofreading and correction. Ecole doctorale Complexité du Vivant (ED 515, Sorbonne Université) and Agence pour la Recherche contre le Cancer for PhD funding of N.R.G. Agence Nationale de la Recherche (ANR-18-CE11-0003; ANR-18-CE11-0003-04; ANR-22-CE12-0004-02) for funding the project. Ecole doctorale IP Paris (ED 626, Institut Polytechnique de Paris) and Fondation ARC pour la Recherche sur le Cancer for PhD funding of I.B.B.

## Author contribution statement

H.L.H, C.S and M.G conceived the project. N.R.G, H.L.H and M.G designed the experiments. N.R.G cloned and purified most proteins and performed *in vitro* pulldowns with the help of J.D and E.A. I.B.B cloned and purified Upf2-Ct and HisZZ-Nmd4. C.G.P cloned GST-Dcp1/Dcp2-L2. M.G performed and analyzed AlphaFold3 simulations, sequence alignments and fluorescence anisotropy assays. H.L.H performed EMSA assays. N.R.G, M.G and H.L.H analyzed the data. N.R.G, H.L.H and M.G wrote the paper with the input of all other authors.

## Reporting summary

Further information on research design is available in the Nature Research Reporting Summary linked to this article.

## Data availability

All data generated or analyzed during this study is included in this published article and its Supplementary Information files. Plasmids generated used in this study are available from the corresponding author upon reasonable request.

