## Supplementary figures for "Mapping interactions within the NMD decapping complex reveals how Upf1 recruits Dcp2 onto mRNA targets"

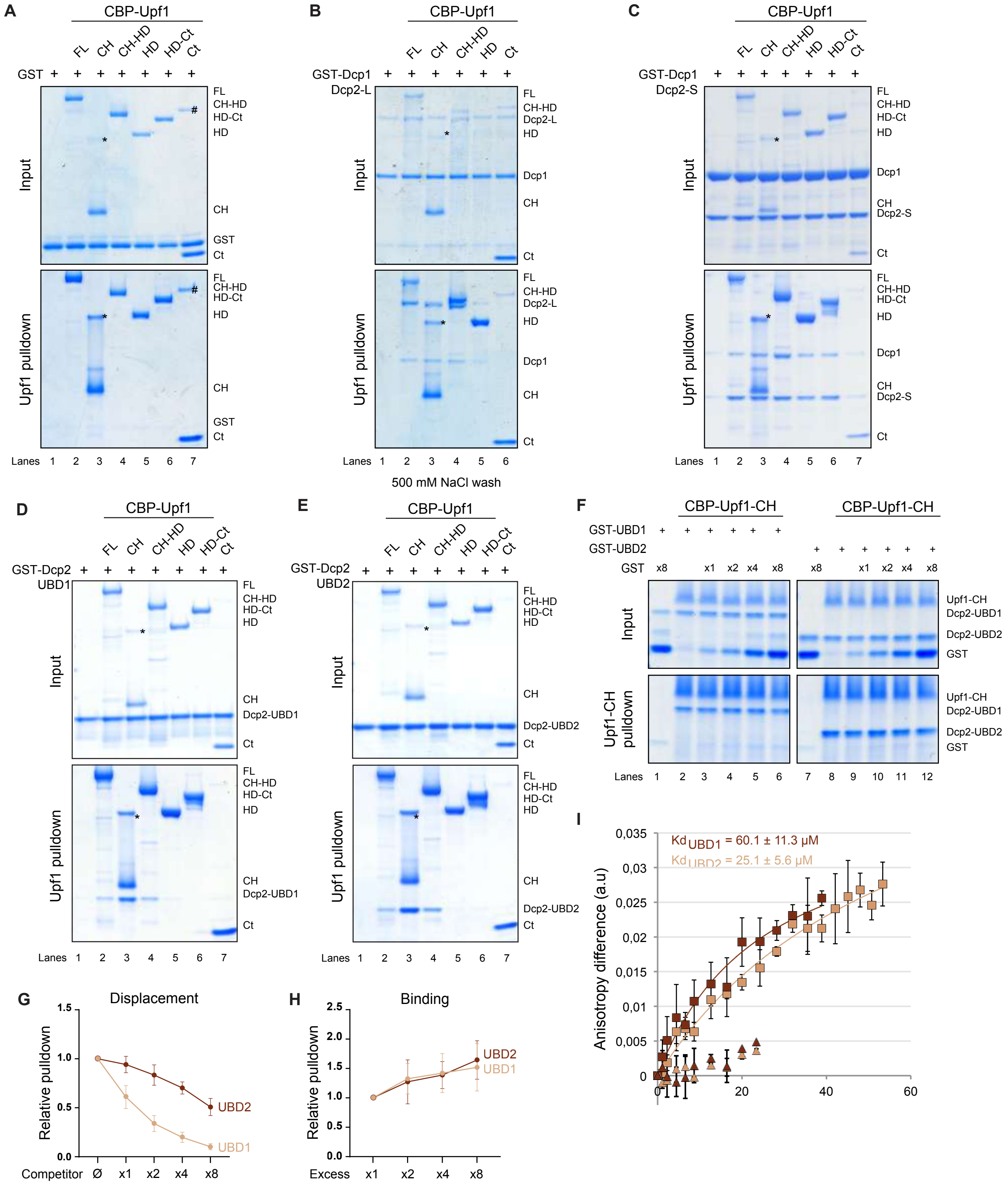


### Figure Supplementary 1

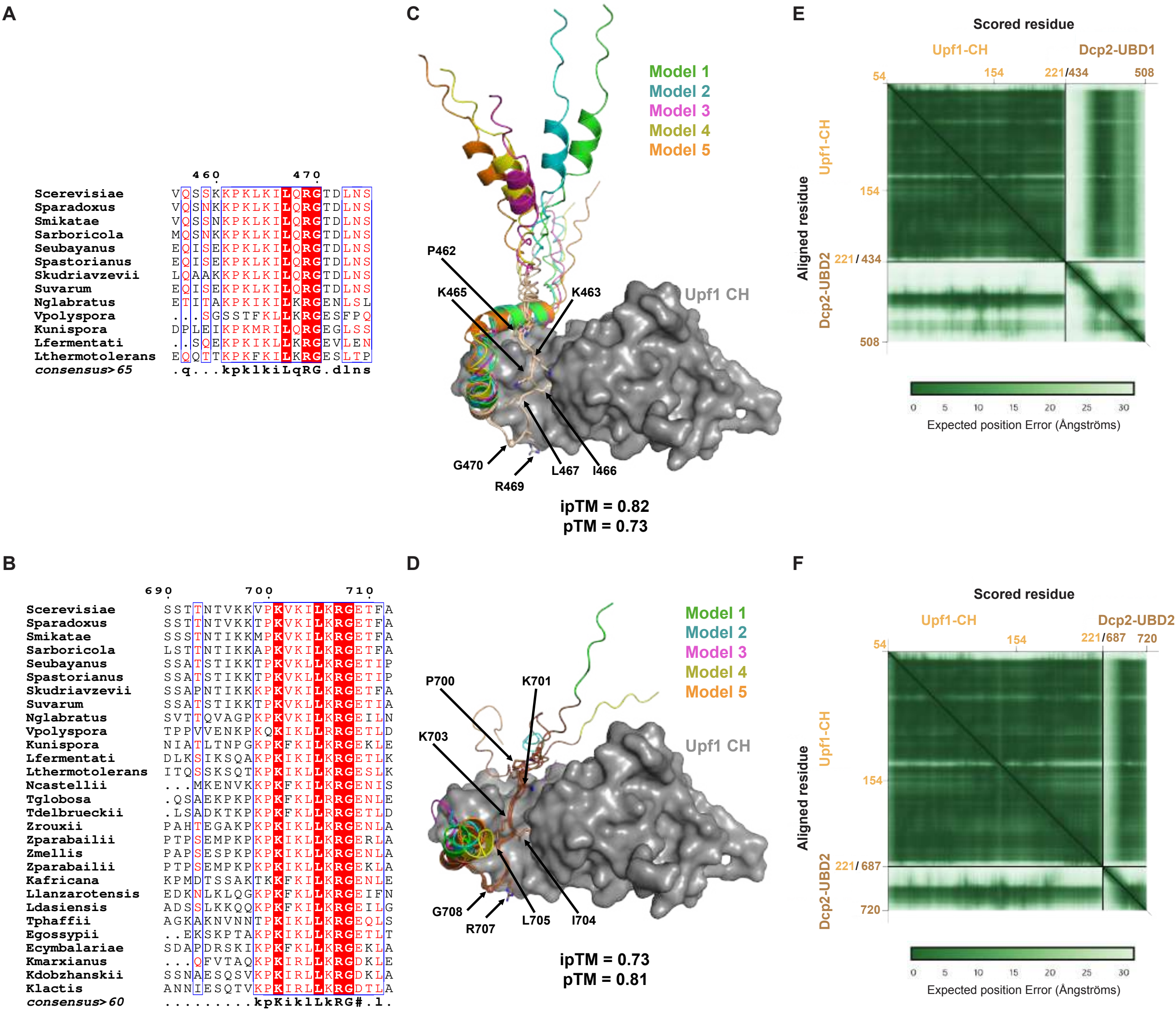


### Figure Supplementary 2

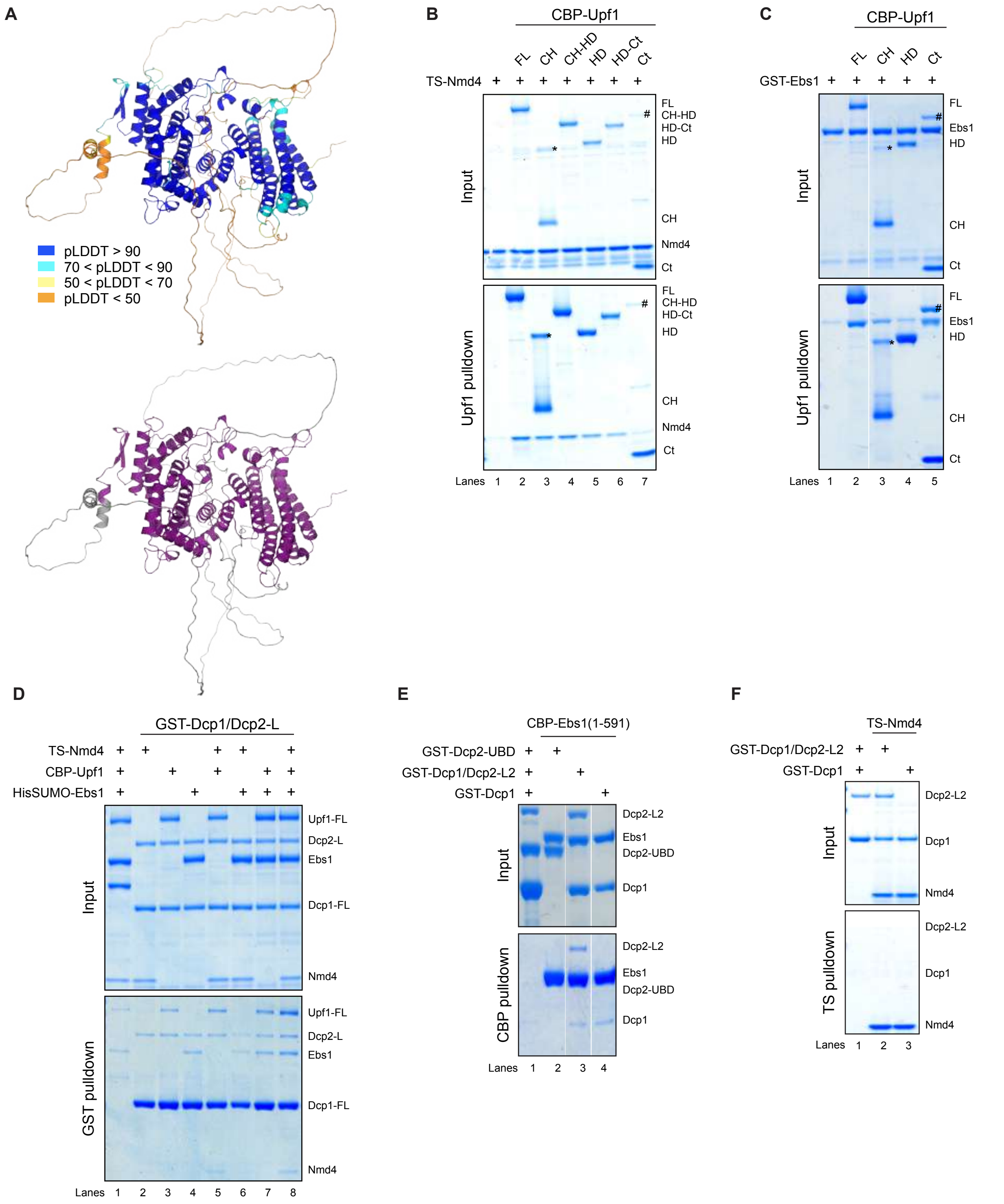


### Figure Supplementary 3

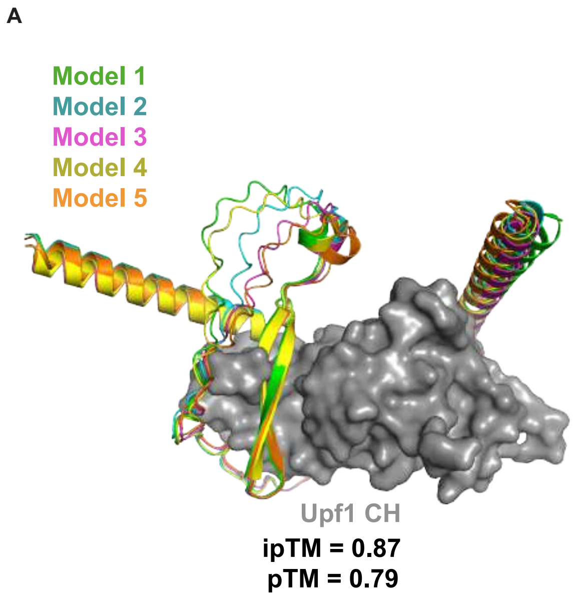


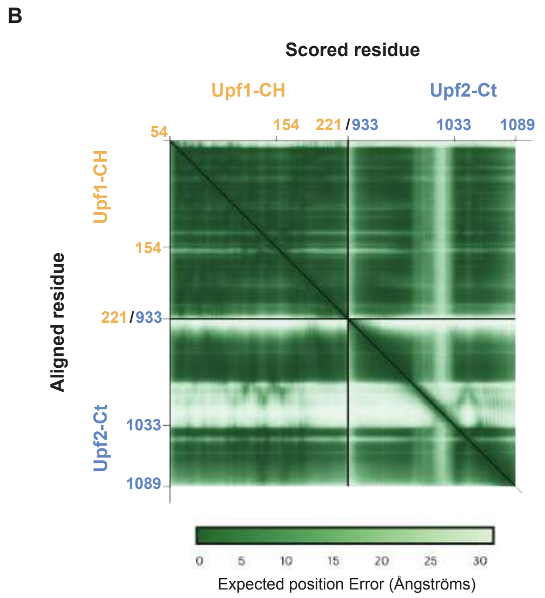


Figure Supplementary 4

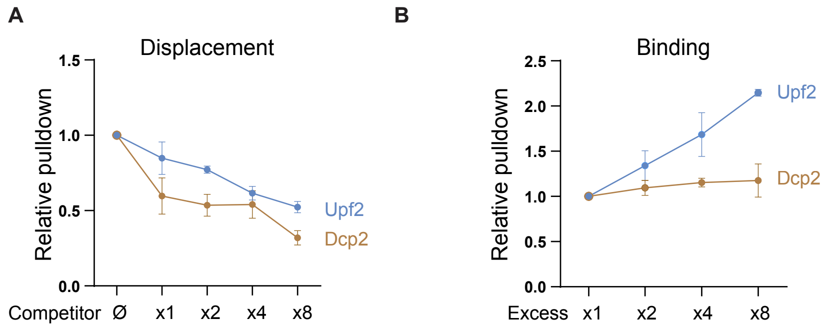


Figure Supplementary 5

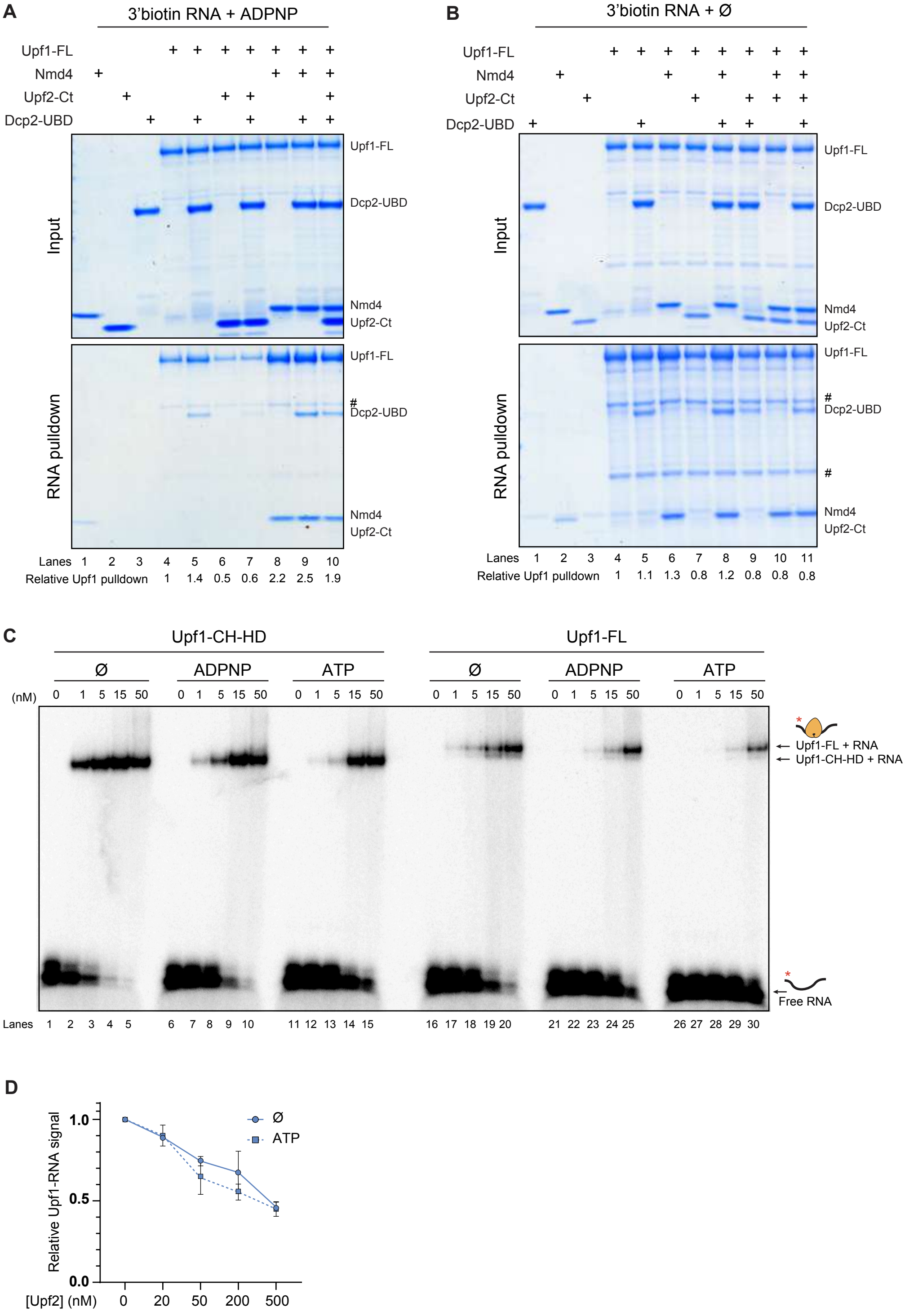


Figure Supplementary 6
